# Microglia undergo rapid phenotypic transformation in acute brain slices but remain essential for neuronal synchrony

**DOI:** 10.1101/2022.04.12.487998

**Authors:** Péter Berki, Csaba Cserép, Balázs Pósfai, Eszter Szabadits, Zsuzsanna Környei, Anna Kellermayer, Miklós Nyerges, Xiaofei Wei, Istvan Mody, Kunihiko Araki, Heinz Beck, Kaikai He, Ya Wang, Zhaofa Wu, Miao Jing, Yulong Li, Attila I. Gulyás, Ádám Dénes

**Author notes:** These authors share first authorship on this work (Péter Berki, Csaba Cserép).

## Abstract

Acute brain slices represent a “workhorse” model for studying the central nervous system (CNS) from nanoscale events to complex circuits. While slice preparation inherently involves tissue injury, it is unclear how microglia, the main immune cells and damage sensors of the CNS shape tissue integrity *ex vivo*. To this end, we have studied the mechanisms of microglial phenotype changes and contribution to neuronal network organisation and functioning in acute brain slices. Using a novel ATP- reporter mouse line and microglia reporter mice, we show that acute slice preparation induces rapid, P2Y12 receptor (P2Y12R) dependent dislocation and migration of microglia, paralleled with marked morphological transformations driven by early ATP surges and subsequent ATP flashes. Gradual depolarization of microglia is associated with the downregulation of purinergic P2Y12R and time-dependent changes of microglia-neuron interactions, paralleled by altered numbers of excitatory and inhibitory synapses. Importantly, functional microglia not only modulate synapse sprouting, but the absence of microglia or microglial P2Y12R markedly diminishes the incidence, amplitude, and frequency of sharp wave-ripple activity in hippocampal slices. Collectively, our data suggest that microglia are inherent modulators of complex neuronal networks, and their specific actions are indispensable to maintain neuronal network integrity and activity *ex vivo.* These findings could facilitate new lines of research resulting in improved *ex vivo* methodologies and a better understanding of microglia-neuron interactions both in physiological and pathological conditions.

## Introduction

Since its first applications several decades ago (Andersen, 1981; Yamamoto & McIlwain, 1966), the acute brain slice preparation technique has become a key tool in the field of neuroscience and has extensively contributed to our understanding of cellular physiology. As the methodology of slice preparation went through numerous iterations over the years (Brahma et al., 2000; Richerson & Messer, 1995; Ting et al., 2014), improvements have led to acute slices even capable of producing spontaneous oscillations, similar to those observed *in vivo* (Hájos et al., 2013; Mann et al., 2005; Perumal et al., 2021; C. Wu et al., 2005). Meanwhile, *ex vivo* studies of non-neuronal cell types such as astrocytes and microglia have also been emerging (Kettenmann et al., 2011; Verkhratsky & Nedergaard, 2018), alongside with increasing focus on bidirectional communication between neurons and glial cells (Eyo & Wu, 2013; Pannasch & Rouach, 2013). Microglia are the primary immune cells of the CNS parenchyma, with essential roles beyond their immune function under both physiological and pathological conditions (Masuda et al., 2020; Prinz et al., 2019). Because microglia are heavily involved in brain development and maintenance of neuronal populations (Kierdorf & Prinz, 2017; Thion et al., 2018), while their functional alterations are linked to a wide range of human diseases (Salter & Stevens, 2017; Song & Colonna, 2018; Wang & Colonna, 2019), interest in understanding microglial function has substantially increased over the last decade. As such, acute slice preparations also proved to be an essential tool for the investigation of microglia, with better preservation of microglial phenotype resembling their physiological states than the *in vitro* culture system (Bennett et al., 2018; Bohlen et al., 2017; Butovsky et al., 2014; Gosselin et al., 2017; Izquierdo et al., 2019), based on robust transcriptomic, proteomic and functional evidence (Boucsein et al., 2000, 2003; Butovsky et al., 2014; Färber & Kettenmann, 2005; Hellwig et al., 2013; Kettenmann et al., 2011; Melief et al., 2012; Schilling & Eder, 2007a; Schmid et al., 2009). However, it is also acknowledged that microglia can become reactive in slices and transform to an amoeboid phenotype (Haynes et al., 2006; Petersen & Dailey, 2004; Stence et al., 2001). Importantly, while the acute slice technique has contributed to understand microglial physiology (Boucsein et al., 2000; Färber & Kettenmann, 2005; Kettenmann et al., 2011; Schilling et al., 2000; Schilling & Eder, 2007b), the impact of methodological practices, relevant time frames, mechanisms influencing microglial states upon slice preparation, and most importantly, the impact of microglial function on electrophysiological measurements have remained vaguely characterized. Despite the known sensitivity of microglia to even subtle changes in their microenvironment (Hirbec et al., 2019; Masuda et al., 2020) mechanistic data from the first few minutes to hours after slice preparation (a relevant time frame for most studies) with appropriate temporal resolution to assess microglial states are not available at present. Thus, it is currently unclear how time-dependent changes of microglia may influence complex neuronal circuits and their reorganization in acute slice preparation. To this end, we set out to investigate microglial phenotype changes and their impact on neuronal networks simultaneously, in slice preparations capable of producing spontaneous network activity by using an experimentally relevant timeframe. For these studies, we made use of established protocols and the extensive expertise in slice preparation available at three independent research laboratories which pioneered the preparation of high quality brain slices and introduced technical developments enabling the first investigation of complex network oscillations (Ferando & Mody, 2015; Hájos et al., 2009; Hájos & Mody, 2009; Remy et al., 2003). Our results suggest that microglia maintain fundamental features of neuronal networks in acute slice preparations, which is remarkably influenced by microglial phenotype changes due to slice preparation-evoked, acute-and sustained ATP release. These observations may be of importance for virtually all acute brain slice studies investigating either neuronal function or microglial responses.

## Results

### Microglia gradually migrate towards the surface of acute slice preparations

Microglia are well known to react rapidly to injury or tissue disturbance in the brain parenchyma (Davalos et al., 2005; Nimmerjahn et al., 2005). To investigate how injury caused by acute slice preparation influences microglial functions at population level, we first tracked microglial cell body and process distribution during a 5-hours incubation in 300 µm thick acute hippocampal slices from CX3CR1^+/GFP^ microglia reporter mice (p.n.:∼35 days). We used a strictly controlled preparation and incubation procedure across all measurements (see details in Methods), optimized for studying spontaneously occurring sharp wave-ripple activity (SWR) (Hájos et al., 2013; Schlingloff et al., 2014). To monitor time dependent changes, slices were immersion-fixed at different timepoints after cutting (Fig 1A). Subsequently, preparations were re-sectioned and mounted onto glass plates for analysis (Fig 1B). Only the native microglial GFP signal was imaged via confocal laser-scanning microscopy (Fig 1C). We found that translocation of microglial cell bodies occurs rapidly after slice preparation with most extensive changes in the top region (∼40 µm) of slices (Fig 1D). Here, the density of cell bodies gradually increased by 75% throughout the 5 hours of the incubation (Mann-Whitney, p<0.01), and reached significance at 42% (p<0.05) as early as 2 hours of incubation (Fig 1E; blue). Top region of slices showed a 42% higher density of cell bodies when compared to the bottom region (Mann-Whitney, p<0.01) after 2 hours of incubation, a relevant timepoint for most electrophysiological measurements (ref, Fig 1E, black statistical indication). We calculated the lowest distance of required cell displacement to reach the 5 hours distribution starting from the 0-minute distribution (Fig 1F). Our results show that the translocation of cell bodies is a composite of two effects: a stronger effect acting in the direction of the top surface and a smaller effect acting towards the top and bottom cut surfaces away from the middle of the slices. The total number of microglia located in whole cross sections did not differ significantly during the experiment (43.07 cells/grid at 0 min, 41.50 cells/grid at 5 hours; p=0.788), suggesting that microglia loss does not contribute to the observed changes in cell distribution.

**Figure 1.**
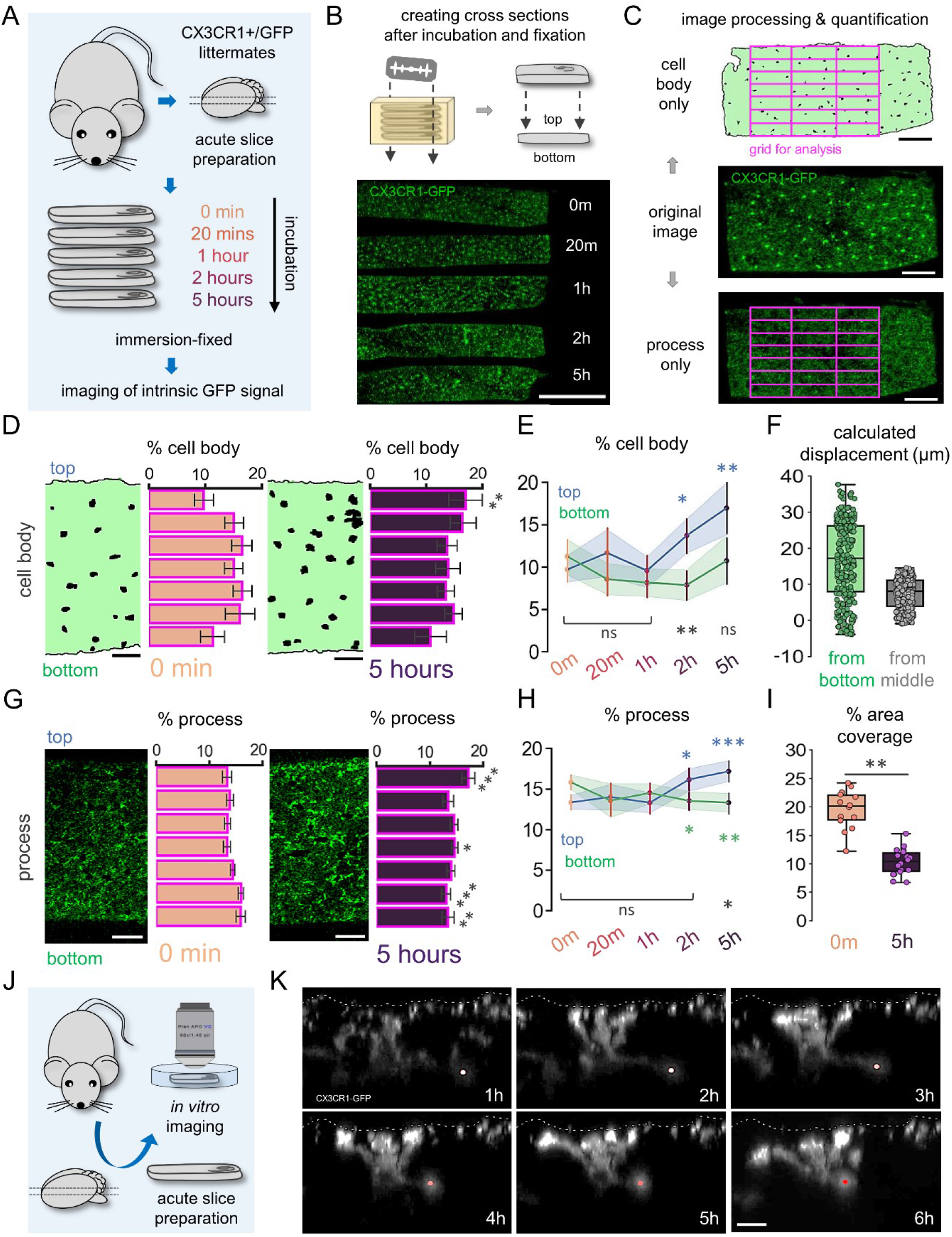
Microglia gradually migrate towards the surface of acute slice preparations. A. Schematic representation of the experiment. CX3CR1^+/GFP^ littermates (N=3; p.n.: ∼35 days) were used to create acute hippocampal slice preparations and placed into an interface-type incubation chamber for recovery. Slices were immersion-fixed immediately (0 minute) or after 20 minutes, 1 hour, 2 hours or 5- hours of incubation. B. Cross sections were made from slice preparations that were fixed at different timepoints (top). Maximum intensity projection image shows cross-sections of slices mounted onto glass plates while preserving their top and bottom directionality according to their position in the incubation chamber. Native GFP signal of microglia were imaged via confocal laser-scanning microscopy (bottom, bar: 500 µm). C. Original images were further processed (to contain either cell bodies or processes) and a 7×3 grid (violet) was used consequently for the quantification of cell body or process distributions along the grid layers (bar: 100 µm). D. Representative sections of processed images at 0 minute and 5 hours showing distribution of microglial cell bodies along the top and bottom axis of slices (left, bar: 50 µm). Bar-plots are showing percentages of total cell bodies counted respective to the layers of the analysis grid (right). N=3, 3 slices/animal, mean ± SEM, independent t-test, **: p<0.01. E. Line-plots representing measured changes of microglial cell body percentages across different timepoints within the top (blue) and bottom (green) layers of acute slice preparations. N=3, 3 slices/animal, median ± SEM, Mann-Whitney, ns: not significant, *: p<0.05, **: p<0.01. Blue statistical indications show significant changes compared to 0-minute timepoint, black statistical indications compare top and bottom means at each timepoint. F. Cell body translocation quantified as minimum required displacement (µm) towards the top measured from bottom (green) or middle (grey). N=3, p.n.: ∼35 days; 3 slices/animal. G. Same as in D) respective to microglial process fluorescence intensity calculated for each layer in the grid, mean ± SEM, Mann-Whitney, **: p<0.01. H. Same as in E) respective to microglial process fluorescence intensity. N=3, p.n.: ∼35 days; 3 slices/animal; mean ± SEM, independent t-test, ns: not significant, *: p<0.05, **: p<0.01, ***: p<0.001. I. Percent of area covered by processes at the 0-minute and 5-hours timepoints, Mann-Whitney, **: p<0.01. J. CX3CR1^+/GFP^ mice (p.n.:45-80 days) were used to create acute hippocampal slices and transferred into a recording chamber for confocal or 2P imaging after cutting. K. Extracted timeframes from Supplementary Video 1. Images show the translocation of microglial processes and the cell body towards the surface of slice preparations (white stripped line). Scale bar: 10 µm.

In line with this, the spatial distribution of microglial processes within the slice showed similar, but even more pronounced alterations during the incubation process (Fig 1G). Microglial process density in the top layer increased significantly by 21% (p<0.05) as early as 2 hours of incubation and by 29% (p<0.001) after 5 hours of incubation (Fig 1H, blue). Contrary to microglial cell bodies, process density at the bottom layer showed a significant drop by 15% (p<0.05) already after 2 hours of incubation (Fig 1H, green). Here, we also observed 23% (p<0.05) lower process density at the bottom region of slices compared to top after 5 hours of incubation (Fig 1H; black statistical indication, independent t-test). Importantly, the total percentage of area covered by microglial processes dropped to half between the 0-minute (acute slices fixed immediately after cutting) and 5-hour timepoints (p<0.0001, independent t-test, Fig 1I).

To capture these changes in real time, slice preparations were transferred into a recording chamber for confocal imaging (Fig 1J). The native signal of microglia (CX3CR1^+/GFP^) was continuously imaged for at least 6 hours after slice preparation (Fig 1K, Supplementary Video 1-2). We found that individual cell behaviours correlated with the quantitative measurements above (Fig 1E-F, H-I), with both processes and the cell body of microglia (Fig 1K, coloured dots) expressing directed movement towards the top surface of slices (Fig 1K, white dashed line).

### Microglia undergo rapid, progressive morphological changes in acute slices

Microglial cells are well known to extend their processes towards injury as an early response, followed by marked morphological transformation (Haynes et al., 2006; Petersen & Dailey, 2004; Stence et al., 2001). Decrease in the area covered by microglial processes (Fig 1I) suggested robust changes in microglia morphology taking place in acute slice preparations. This behaviour was also evident in the data gathered by our imaging experiments, where we saw rapid morphological transformation of individual microglial cells (Fig 2A, Supplementary Video 3). To define the spatio-temporal characteristics of these pronounced morphological changes, preparations were immersion-fixed at different timepoints during incubation. In parallel, a group of mice were transcardially perfused to obtain slice preparations from the same brain area as controls (Fig 2B). Confocal images were analysed using unbiased automated morphological analysis (Fig 2C; Heindl et al., 2018).

**Figure 2.**
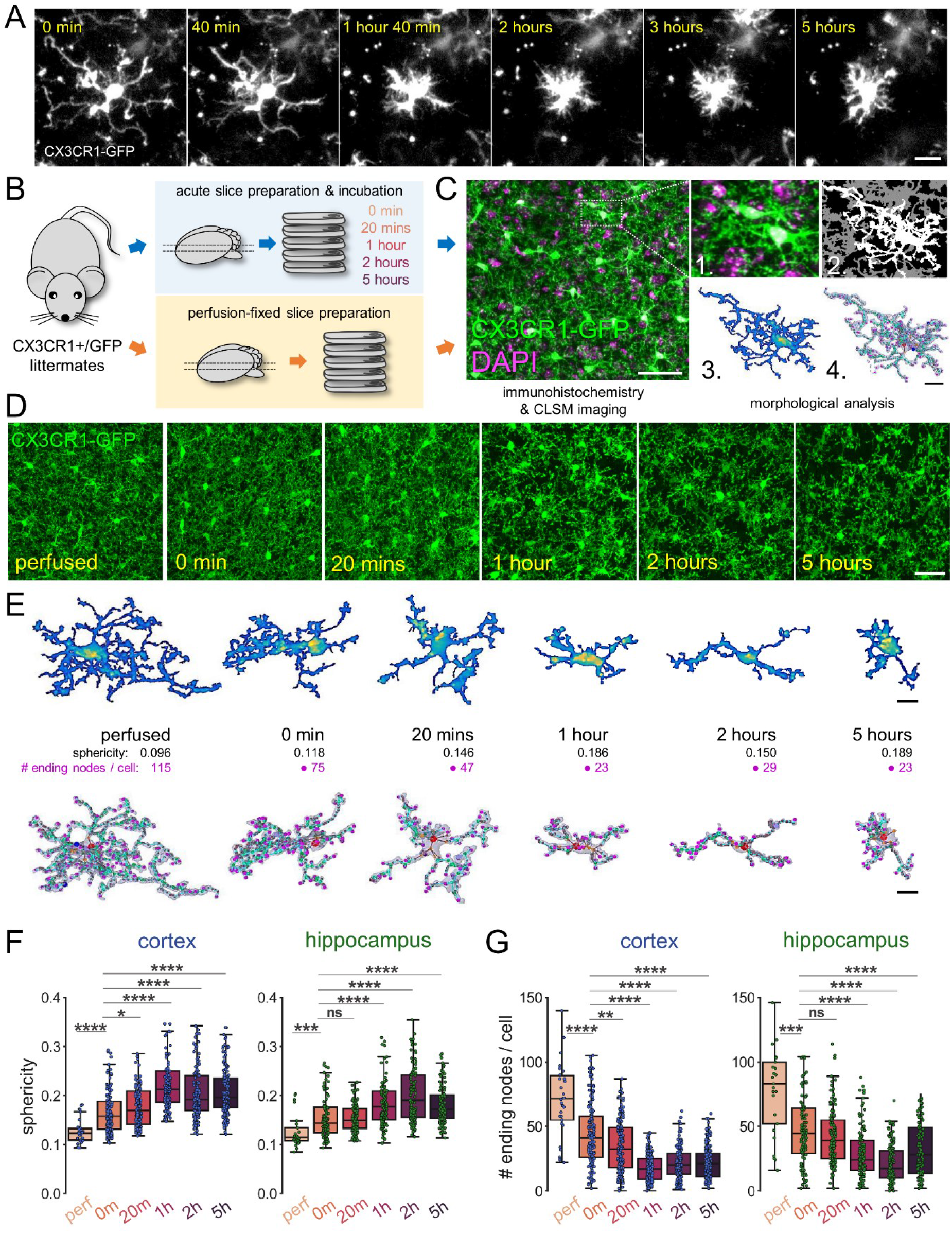
Microglia undergo rapid, progressive morphological changes in acute slices. A. Extracted timeframes from Supplementary Video 3. CX3CR1^+/GFP^ signal of a single microglial cell was captured via confocal imaging at different timepoints after acute slice preparation. Scale bar: 10 µm. B. Schematic representation of experiment. CX3CR1^+/GFP^ littermates (N=8; p.n.: ∼35 days) were used either to create hippocampal slice preparations, which were immersion-fixed before (0 minute) and after 20 minutes, 1 hour, 2 hours, or 5 hours of incubation (blue arrows), or animals were perfusion fixed to obtain slice preparations with the same dimensions as acute slices (red arrows). C. Slices were stained and z-stack images were obtained via confocal laser-scanning microscopy (left, bar: 20 µm). Images were analysed with an automated morphological analysis tool (Heindl et al, 2018) which uses raw z-stacks (1.) to isolate and segment the images to separate microglial cells (2.). Separate cells are further segmented (3.) to cell body (yellow) and processes (blue, purple). Finally, a skeleton is constructed (4.) representing a 3D-model of each individual microglial cell (right, bar: 10 µm). D. Maximum intensity projections of analysed confocal images showing microglial cells in perfused or immersion-fixed slices (bar: 50 µm). E. Top row: individual microglial cells at each timepoint respective to section C) above (yellow: cell body; blue, purple: processes; bar: 5 µm). Bottom row: reconstructed skeletons for each cell depicted in middle row, together with their sphericity values (black) and total number of process endings (violet, cell body: red dot, bar: 5 µm). F. Quantification of extracted morphological features regarding sphericity in cortex (green) and in hippocampus (pink). N=8 animal, p.n.: ∼35 days; Mann-Whitney, ns: not significant, *: p<0.05, **: p<0.01, ***: p<0.001, ****: p<0.0001. G. Same as in E) regarding number of process endings / cell in cortex (green) and in hippocampus (pink). N=8 animal, p.n.: ∼35 days; Mann-Whitney, ns: not significant, *: p<0.05, **: p<0.01, ***: p<0.001, ****: p<0.0001.

**Supplementary Fig 1:**
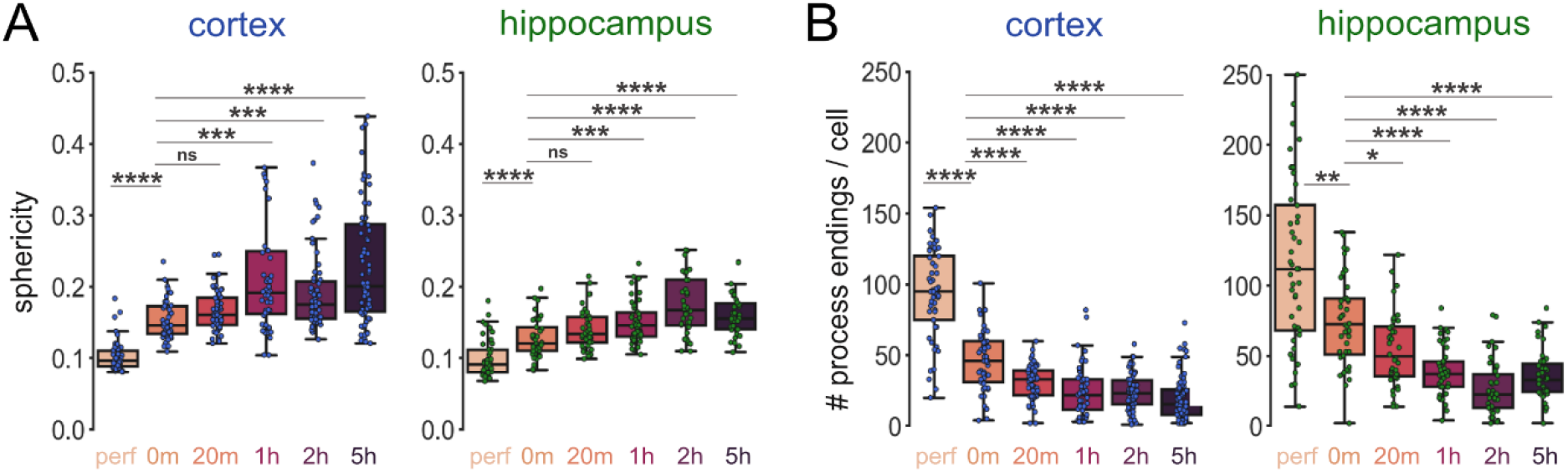
Morphological changes of microglia in acute slice preparations obtained from an older group of mice (p.n.: ∼95 days) A. Quantification of extracted morphological features regarding sphericity in cortex (green) and in hippocampus (pink). CX3CR1^+/GFP^ littermates, N=5 animal, p.n.: ∼95 days; Mann-Whitney, ns: not significant, *: p<0.05, **: p<0.01, ***: p<0.001, ****: p<0.0001. B. Same as in A) regarding number of process endings / cell in cortex (green) and in hippocampus (pink). N=5 animal, p.n.: ∼95 days; Mann-Whitney, ns: not significant, *: p<0.05, **: p<0.01, ***: p<0.001, ****: p<0.0001.

Slice preparation induced drastic morphological differences in microglia. We observed rapid retraction of processes already visible in maximum intensity projection images created from the analysed z-stacks (Fig 2D), as well as on individual cell morphologies and skeleton reconstructions via the analysis tool (Fig 2E; Supplementary Video 4). Sphericity values of individual cells (amoeboid shape and retraction of processes translate to higher sphericity values) increased by ∼50% on average already after 20 minutes of incubation (independent t-test, p<0.0001), and peaked at ∼100% increase (p<0.0001) between 1 and 2 hours of incubation both in regions of the cortex and in the hippocampus (Fig 2F). At the same time, total number of process endings dropped by ∼50% after 20 minutes of incubation (p<0.0001), and this reduction also peaked between 1- and 2 hours of incubation with ∼80% decrease in average (Fig 2G, p<0.0001). We also conducted this experiment in an older group of animals (p.n.: ∼95) and observed the same extensive changes concerning both cortical and hippocampal regions (Supplementary Fig 1). Based on these observations, we concluded that microglia showed rapid morphological changes in acute slice preparations.

### Migration and rapid phenotype changes of microglia during the incubation process do not depend on cutting techniques

To investigate whether different slice cutting techniques could influence the extent or temporal course of microglial phenotype changes during the incubation process, we next compared acute slice samples across three independent laboratories (Lab #1, Lab #2, Lab #3) using identical fixation protocol and timing for serial preparations, but left native, well-established slice cutting techniques intact by each laboratory. Briefly: Lab#1 used an ice-cold, sucrose-based cutting solution, and slices were incubated in an interface-type chamber filled with carbonated standard ACSF. Lab #2 used an ice-cold, N-methyl-D-glucamine (NMDG)-based cutting solution, and slices were incubated in a low Na^+^, sucrose-based ACSF while floating in a Styrofoam “boat” with a netting at the bottom on the surface of an incubation chamber; Lab #3 used an ice-cold, sucrose-based ACSF solution, and slices were incubated in a storage chamber filled with carbonated standard ACSF (for details see: Methods). Of note, we also tested slice preparation using a room-temperature cutting solution, which did not alter the course of microglial changes (data not shown). In all laboratories, slices were immersion-fixed in 4% PFA. Then, histological processing, imaging and analysis were performed in a uniform and blinded manner at the IEM (Fig 3A-C). Confirming our earlier results, both the sphericity of microglial cells (Fig 3D) and the number of ending nodes (Fig 3E) showed closely identical changes, similarly to microglial cell body migration and process movements (Fig 3F) suggesting that the observed changes are general, common characteristics of different acute slice preparation techniques.

**Figure 3.**
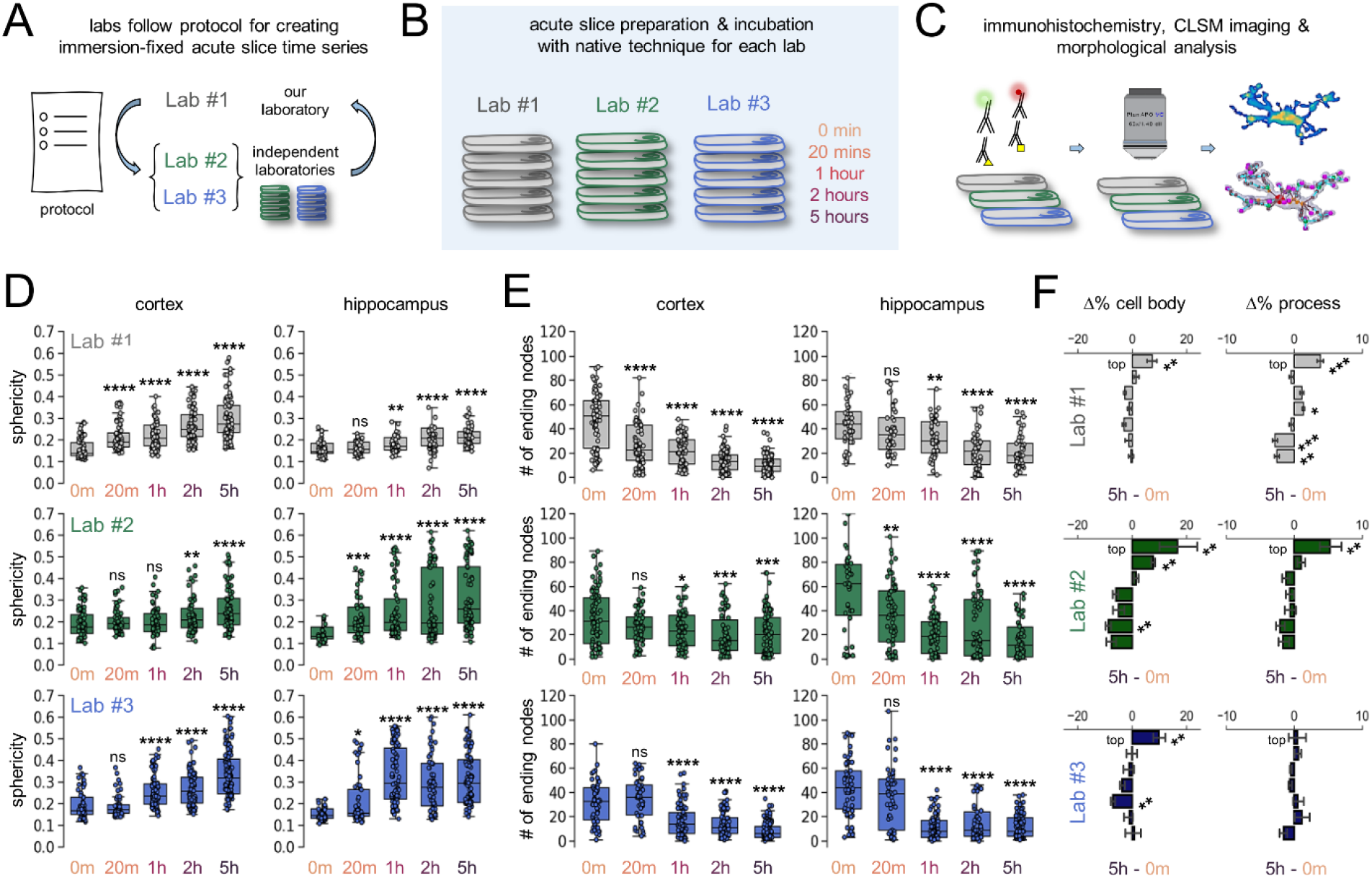
Migration and rapid phenotype changes of microglia during the incubation process do not depend on cutting techniques. A. Schematic representation of experiment. To compare different laboratories and acute slice preparation techniques, two independent laboratories received a protocol to synchronize acute slice fixation timepoints and fixation methods. B. Independent laboratories were using their own native acute slice preparation method to harvest and fixate slices at specific timepoints according to the protocol. C. Acute slice preparations were sent to our laboratory after fixation and preparation for transport (see: Methods), where they were treated together during immunostaining, imaging, and morphological analysis. D. Quantification of extracted morphological features across different laboratories regarding sphericity in cortex (left) and in hippocampus (right). N=2 animal/lab, p.n.: ∼65 days; Mann-Whitney, ns: not significant, *: p<0.05, **: p<0.01, ***: p<0.001, ****: p<0.0001. E. Same as in D, regarding number of ending nodes/cell in cortex (left) and in hippocampus (right). N=2 animal/lab, p.n.: ∼65 days; Mann-Whitney, ns: not significant, *: p<0.05, **: p<0.01, ***: p<0.001, ****: p<0.0001. F. Bar plots representing measured changes of microglial cell body numbers (left) and area covered by processes (right) in percentages calculated between 0 min and 5 hours across the top and bottom layers of acute slice preparations. N=2 animal/lab, median ± SEM, Mann-Whitney, ns: not significant, *: p<0.05, **: p<0.01, ***: p<0.001.

### Rapid downregulation of P2Y12R is accompanied by gradual microglial depolarization during the incubation process

To further investigate the extent of microglia transformation during the incubation process, we next examined changes in microglial P2Y12 receptor (P2Y12R) levels, a core microglial receptor through which microglia sense and influence neuronal activity and fate (Cserép et al., 2020; Dissing-Olesen et al., 2014; Eyo et al., 2014; Gu et al., 2016; Kato et al., 2016), and which is downregulated upon injury and inflammatory challenges (Haynes et al., 2006; Mildner et al., 2017). Pre-embedding immunofluorescent labelling indicated a marked reduction of P2Y12R labelling intensity already 1 hour after slice preparation (Fig 4A). To quantitatively examine these changes during the whole incubation process, acute slice preparations were immersion-fixed at different timepoints during incubation (Fig 4B), followed by visualization of P2Y12R with a recently developed quantitative post-embedding immunofluorescent labelling technique (Holderith et al., 2020), enabling unbiased assessment of P2Y12R labelling intensity changes (Fig 4C). We quantified P2Y12R labelling intensity at microglial cell bodies (Fig 4D, left), thick processes (Fig 4D, middle) and thin processes (Fig 4D, right), to investigate whether different subcellular compartments are differentially affected. Confirming our immunofluorescent results, we found a consistent, rapid decrease in P2Y12R labelling intensity in all the compartments until 1 hour of incubation, followed by a slight increase towards the 5-hour timepoint (Fig 4E). Loss of P2Y12R was most prominent in thick and thin processes (Fig 4E, middle, right), and less apparent on microglial cell bodies (Fig 4E, left). As such, already after 1 hour of incubation, the labelling intensity of P2Y12R (arbitrary unit, see: Methods) dropped by 25% on thick processes (p<0.01), 29% on thin processes (p<0.001) and 20% (p<0.05) at cell bodies of microglia.

**Figure 4.**
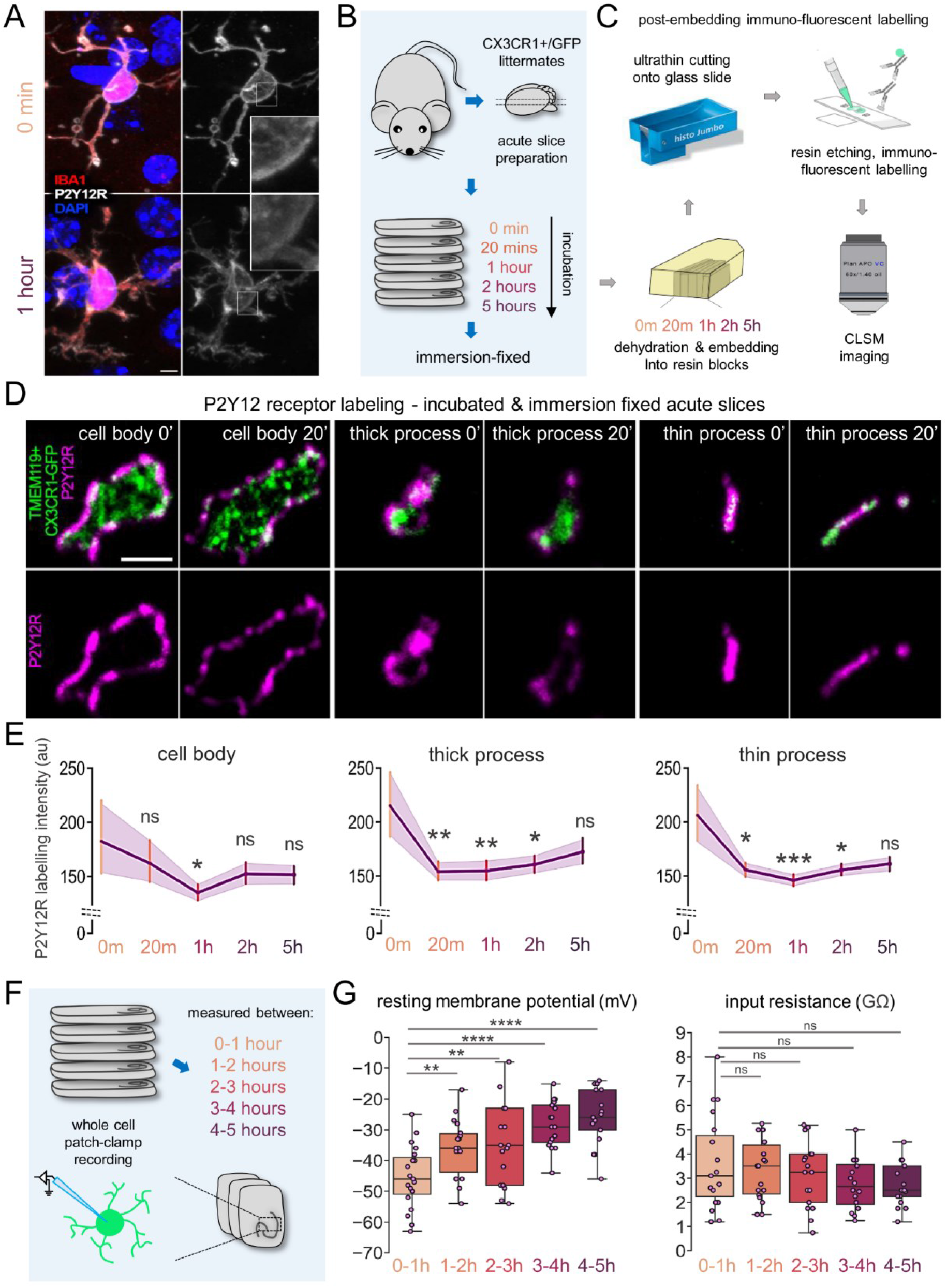
Rapid downregulation of P2Y12R is accompanied by gradual depolarization of microglia during the incubation process. A. Single image planes of multi-channel confocal laser scanning microscopy z-stacks, depicting microglial cells with pre-embedding immunofluorescent labelling (IBA1, red; P2Y12 receptors, white) before (0 min) and after 1 hour of incubation (bar: 3 µm). B. Schematic representation of the experiment. CX3CR1^+/GFP^ littermates (N=3; p.n.: ∼35 days) were used to create acute hippocampal slice preparations and placed into an interface-type incubation chamber for recovery. Slices were immersion-fixed before (0 minute) or after 20 minutes, 1 hour, 2 hours, and 5 hours of incubation. C. After fixation, P2Y12 receptor labelling was performed by a post-embedding immunofluorescent labelling technique: slices were dehydrated and embedded into resin blocks. Subsequently, ultrathin slices were cut onto glass slides and labelled after resin etching. Finally, z-stack images were gathered from preparations via high-resolution confocal laser scanning microscopy. D. Representative images of P2Y12 receptor labelling via the post-embedding technique. Microglial cell bodies (left), thick processes (middle) and thin processes (right) are shown together with the P2Y12R labelling (top row) and P2Y12R labelling only (bottom row) at 0- and 20-minute timepoints (Scale bar: 2 µm). E. Quantification of P2Y12 receptor labelling intensity (arbitrary unit, see: Methods) in microglial cell body (left), thick processes (middle) and thin processes (right.) N=3 animal, p.n.: 35 days; 3 slices/animal; Mann-Whitney, ns: not significant, *: p<0.05, **: p<0.01, ***: p<0.001. F. Schematic representation of the experiment. CX3CR1^+/GFP^ littermates (N=10, p.n.: ∼95 days) were used to create acute slices. Microglial cells were measured in whole cell patch-clamp configuration. Microglial cells were targeted via their intrinsic GFP signal across the hippocampal CA1-CA3 stratum lacunosum-moleculare and stratum radiatum regions. G. Resting membrane potential (left) and input resistance (right) of individual microglial cells at different timepoints of the recovery process. N=10 animal, p.n.: ∼90 days; N=158 cells measured in total; independent t-test, ns: not significant, *: p<0.05, **: p<0.01, ***: p<0.001.

Microglial phenotype changes have been shown to correlate with resting membrane potential, while the tonically active K^+^ channels responsible for maintaining resting membrane potential are known to be potentiated by P2Y12R actions (Madry, Kyrargyri, et al., 2018; Swiatkowski et al., 2016), especially in response to injury when exposure to high ATP/ADP levels. Therefore, rapid P2Y12R downregulation is expected to be accompanied with membrane depolarization. To test this, we performed electrophysiological recordings from microglia in acute slices. As in previous experiments, acute slice preparations were placed into an interface type incubation chamber for recovery and then transferred to a recording chamber at different timepoints during incubation, to measure microglia in whole cell patch-clamp configuration. Targeting of microglial cells was guided by the intrinsic GFP signal across the hippocampal CA1-CA3 stratum lacunosum-moleculare and stratum radiatum regions, and below ∼40 µm measured from the slice surface (Fig 4F). Our results showed that microglial cells became gradually more depolarized throughout the total 5 hours of the incubation process (Fig 4G, left; N=158 cells measured in slices from a total of 10 animals), while we did not observe significant changes in input resistance (Fig 4G, right). Taken together, microglia undergo rapid phenotype changes in acute slice preparations, as characterized by cell body and process translocation (Fig 1, 3), quick morphological shift towards a reactive shape (Fig 2, 3), as well as early P2Y12R downregulation and gradual depolarization of resting membrane potential (Fig 4).

### Immediate release of ATP and sustained ATP fluctuations contribute to microglial phenotype changes in a P2Y12R dependent manner

Since injury-induced ATP markedly affects microglial phenotypes and recruitment via P2Y12R (Davalos et al., 2005; Haynes et al., 2006), we set out to examine extracellular ATP levels in acute brain slices by generating a novel mouse strain expressing a sensitive, genetically-encoded fluorescent ATP sensor (GRABATP) in VGlut1 positive neurons (Z. Wu et al., 2022). In order to capture time-dependent extracellular ATP dynamics, we recorded 10 min long videos via two-photon imaging, as well as confocal Z-stacks from both regions of cortex and hippocampus in each hour after the slice preparation (Fig 5A). We observed high fluorescent signals, representing extracellular ATP levels both in the hippocampal CA3 region and in cortical layers shortly after cutting (Fig 5B), which was most intense near the slice surface, as revealed by Z stack acquisitions. The mean ATP signal rapidly declined in the first 10 minutes of imaging (Fig 5B-C; Suppl. Fig 2A), corresponding to the 10-20 min interval upon slice preparation (see: Fig.5.A and Methods). Then on, ATP levels showed a slow, gradual decrease (Fig 5D, F; Suppl. Fig 2B; Supplementary Video 5) with ATP gradient maintained through the 0-100 µm depth of the tissue for at least 5 hours (Suppl Fig 2C). While ATP release from brain slices generally decreased over time, we also observed intense, focal ATP flashes throughout the 5 hours period after slice preparation (Supplementary Video 5; Fig 5E). ATP flash incidence in acute slices was 12.3±2.2 (hippocampus) and 25.4±5.2 (cortex) during the 10 min imaging periods, resembling systemic inflammation-induced flashing activity *in vivo* (Z. Wu et al., 2022) (Fig.5.H, Suppl. Fig 3A). While we detected more frequent and intense ATP flashes in cortical regions of acute slices than in the hippocampus (Fig 5 G-H), all other characteristics (ATP flash area, duration, rise time, decay time) were quite uniform (Suppl. Fig 3B), in agreement with earlier observations (Z. Wu et al., 2022). It should be noted, however, that the observed intensity differences across different depth of slices may also be affected by the heterogenous sensor expression.

**Figure 5.**
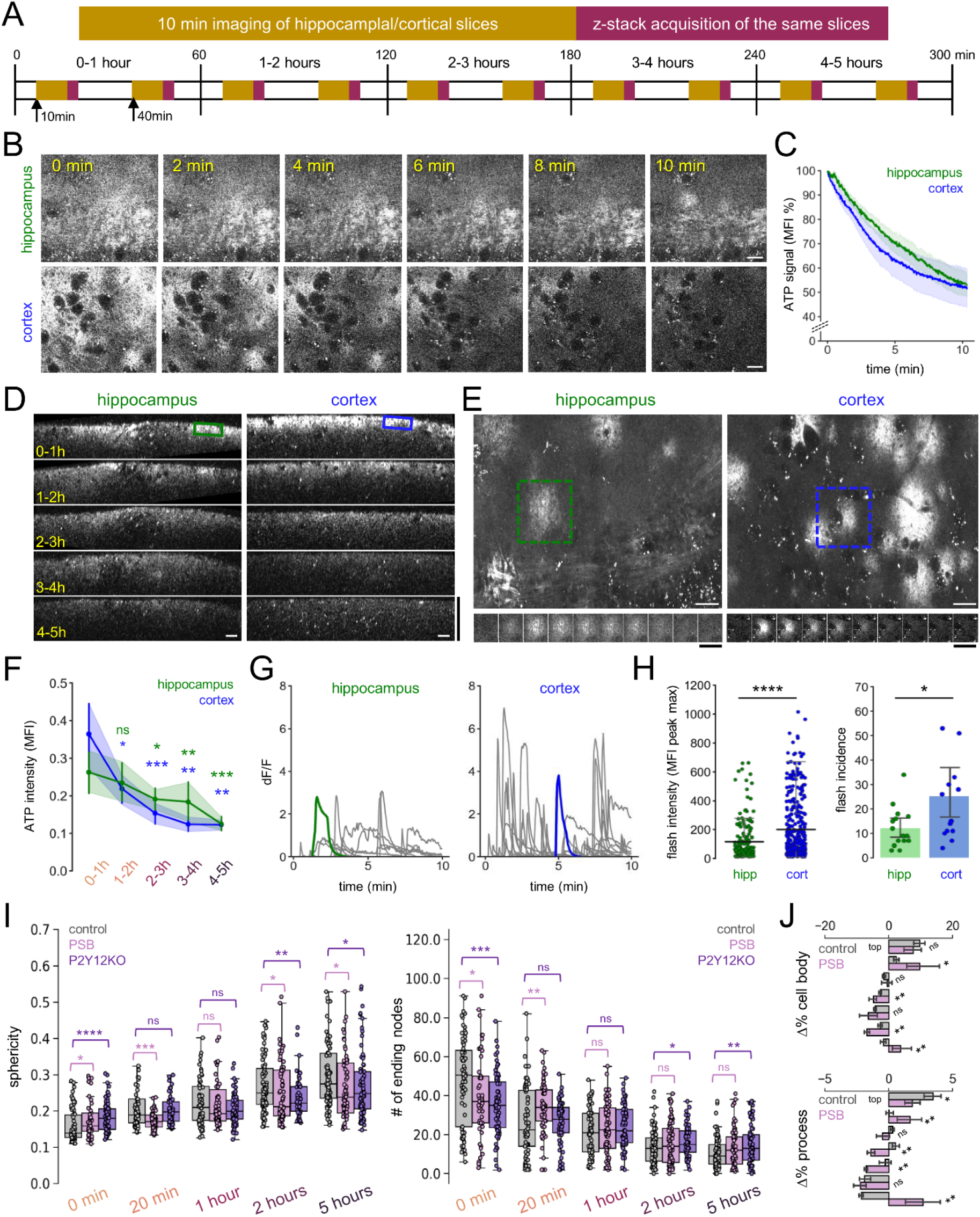
Immediate release of ATP and sustained ATP fluctuations contribute to microglial phenotype changes in a P2Y12R dependent manner. A. Outline of the two-photon imaging experiment performed on acute brain slices. Slices were prepared from mice expressing the ATP sensor (GRABATP on the cell surface of VGlut1-positive neurons to measure changes in extracellular ATP levels. 10 min-long imaging sessions were followed by Z-stack acquisition after slice cutting, for up to 5 hours. Altogether, 3*5 hippocampal (CA3) and 3*5 cortical (layer 2-3) data sets were collected from n=3 mice 1 hour apart. B. Representative images of the video sets taken at the earliest manageable timepoint after slice cutting. Scale bar: 20 µm. C. MFI data indicating a continuous and significant decrease of the extracellular ATP signal in regions with no flashing ATP activity (Two-way repeated measures ANOVA, F(1.221, 23.20)=71.63, ****P<0.0001). A gradual, slow decrease of ATP signal was observed in subsequent timepoints (see Suppl. Fig 2A). D. Representative side-views of Z stacks show marked ATP production next to the slice surface (0-25µm) showing a time-dependent decrease of the signal towards the 5 hours’ timepoint. Horizontal scale bar: 20 µm, vertical scale bar: 125 µm. E. Maximum intensity projection images of representative videos show the occurrence of ATP flashes. Below the images the timeline of two individual ATP flashes are shown (10 sec/image). White scale bar: 20 µm, black scale bar: 10 sec. Quantification of the MFI measurements shown in D, as ATP levels are following a significant, time-dependent decrease towards the 5 hours’ timepoint. Two-way repeated measures ANOVA with Tukey’s multiple comparison test; data are compared to 0-1h values for either cortex or hippocampus: *:p<0.05, **:p<0.01, ***:p<0.001 F. Corresponding dF/F graphs depict flashing activity at the same fields as shown in E. G. The intensity of the ATP flashes is significantly higher in cortical slices (n=360) than in hippocampal ones (n=181). Mann-Whitney test, ****p<0.0001. The incidence of flashes is higher in cortical slices. The number of ATP flash/10min imaging/slice are shown. Unpaired t-test, P=0.0258. H. Quantification of extracted morphological features showing significant differences between control and P2Y12KO or PSB treated acute slices regarding the sphericity (left) and the number of ending nodes/cell (right). N=2 animal/lab, p.n.: ∼65 days; Mann-Whitney, ns: not significant, *: p<0.05, **: p<0.01, ***: p<0.001, ****: p<0.0001. I. Bar plots showing the differences between control and PSB treated acute slices. Changes of microglial cell body percentages (top) and area covered by processes (bottom) are calculated between 0 min and the 5 hours across the top and bottom layers of acute slice preparations. N=2 animal/lab, median ± SEM, Mann-Whitney, ns: not significant, *: p<0.05, **: p<0.01.

**Supplementary Figure 2.**
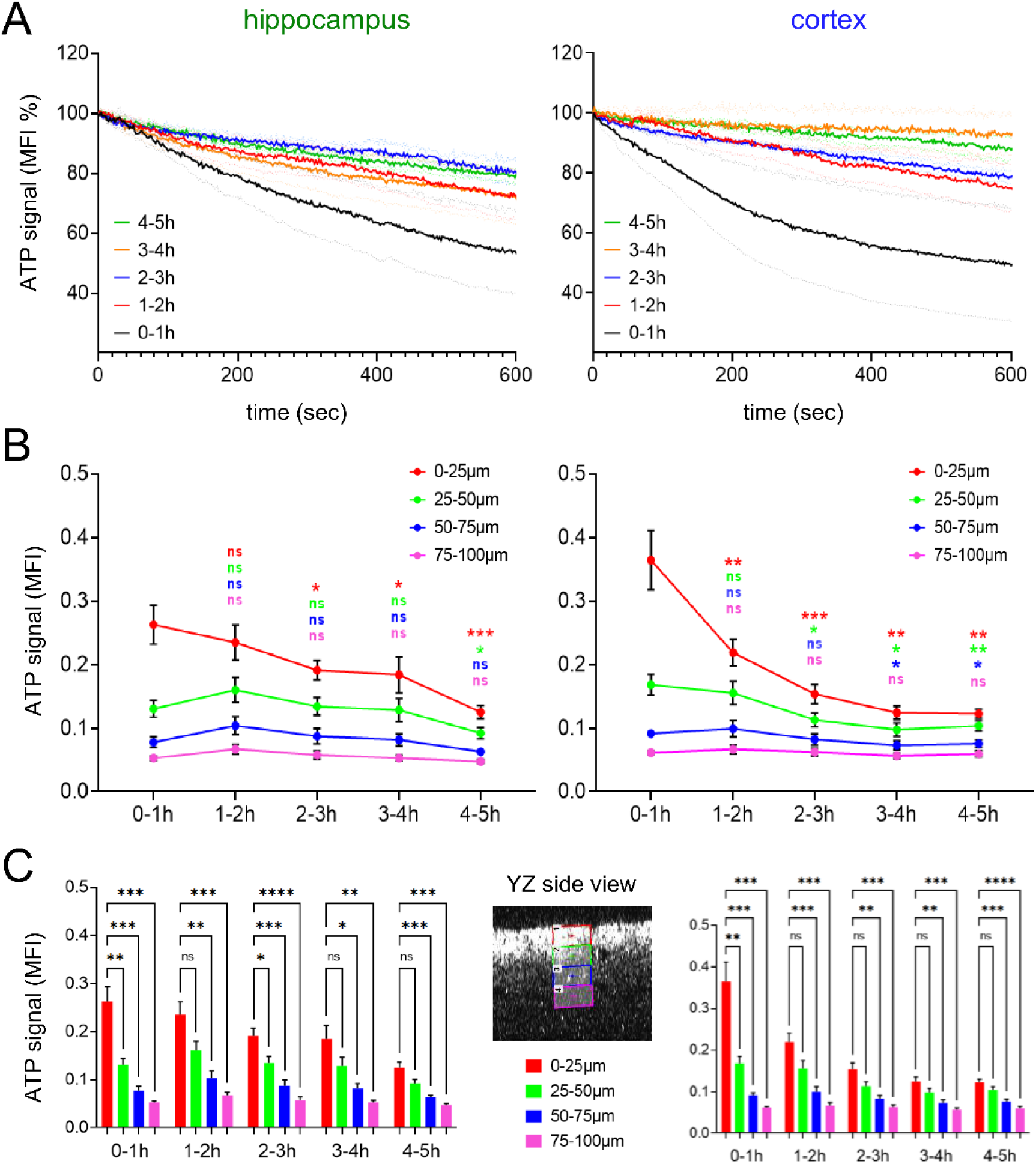
Characteristics of extracellular ATP signal changes in acute brain slices. A. MFI data collected from hippocampal or cortical slices imaged 1-2h (red), 2-3h (blue), 3-4h (orange) or 4-5 (green) hours after slice preparation show significantly slower ATP reduction, than those from the earliest timepoints (0-1h, black). Note, that data were collected from regions of interests where no apparent flashing ATP activity was observed. Two-way ANOVA, ****P<0.0001 for all (red, blue, orange, green) compared to 0-1h (black). B. MFI data obtained from Z stacks. ATP signal intensity and subsequent reduction over time is most apparent next to the slice surface (0-25, 25-50um) compared to deeper layers (75-100um). Two-way repeated measures ANOVA, Dunnett’s multiple comparison test. C. Gradient in ATP signal intensity from the slice surface to the ∼100 µm depth of the slice is maintained for at least 5 hours after slice preparation. Two-way repeated measures ANOVA, Dunnett’s multiple comparison test.

**Supplementary Figure 3.**
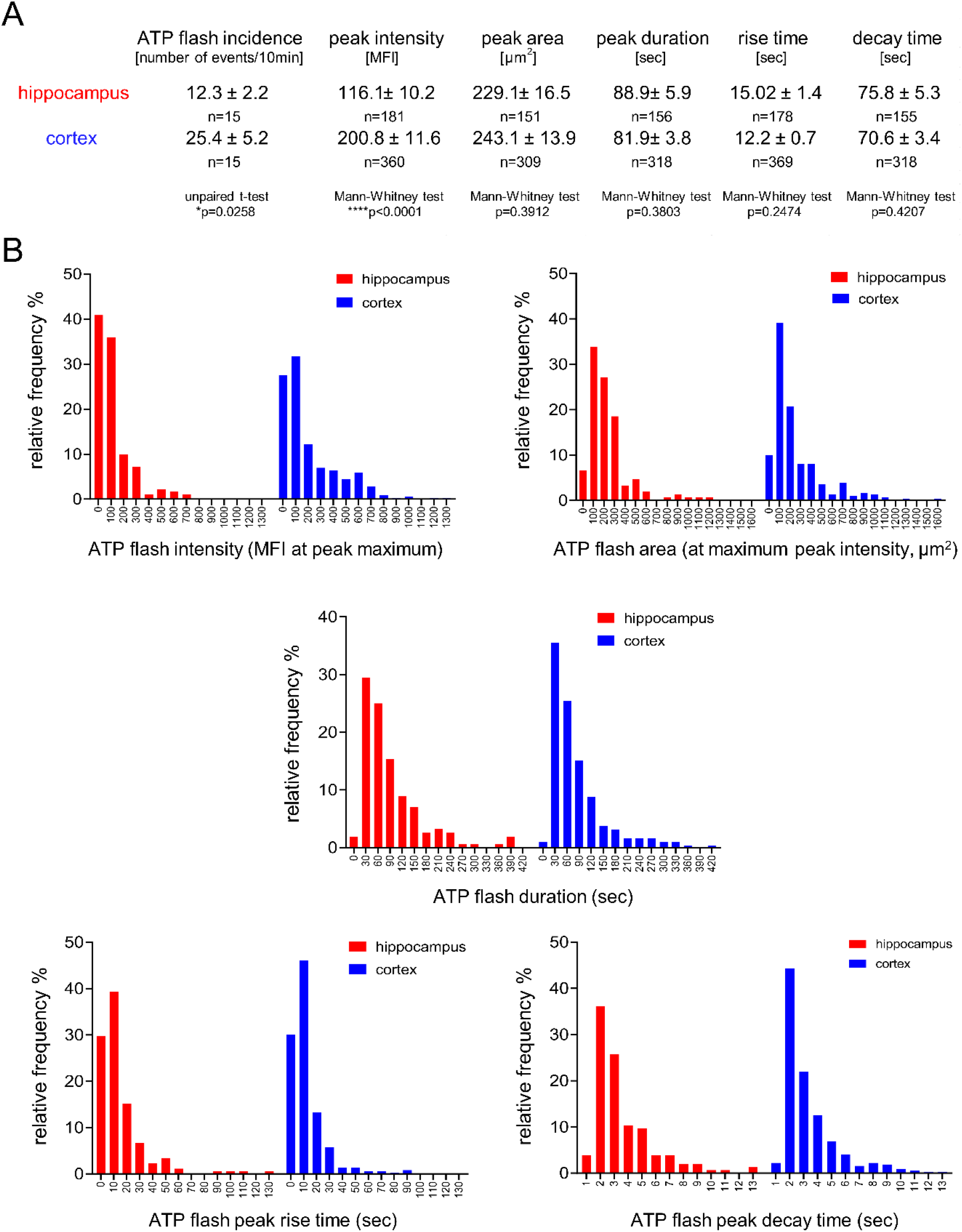
Properties of ATP flash events in acute slices. A. Statistics of ATP flash properties. B. Histograms show data distribution of flash intensity, area, duration, rise time and decay time.

In order to directly test whether purinergic signaling would play a role in the observed migration and morphological changes of microglia, we prepared samples from wild-type, P2Y12R-KO and PSB0739 (PSB, a selective P2Y12R-inhibitor) treated wild-type mice, where PSB (10 µM) was present both in the cutting and incubating solutions during slice preparation. Time series of slices were prepared and fixed as previously. We found that the time course of the changes, including microglial sphericity and number of ending nodes, as well as dislocation and process redistribution of microglia were disrupted significantly by the absence or inhibition of P2Y12Rs (Fig 5I and J), (Fig 5J), suggesting that microglial phenotypic changes during slice incubation are influenced by extracellular ATP (ADP) via P2Y12R.

### Rapid changes of microglia-neuron interactions and microglia-dependent synaptic sprouting characterize acute slices

Next, we investigated whether changes in microglial phenotype parallel altered microglia-neuron interactions in acute slice preparations. To this end, we looked at microglial contacts with synaptic elements and neuronal cell bodies (Cserép et al., 2020), in order to examine how the number of these contacts might change during the incubation process. The native GFP-signal of microglia was enhanced by anti-GFP labelling, while neuronal cell bodies were identified via Kv2.1 labelling. Synaptic elements were further identified via VGLUT1-Homer1 and VGAT-Gephyrin co-localization (Fig 6A,C). We found that the total percentage of neuronal soma contacted by microglial processes gradually decreased throughout the incubation process and dropped by ∼23% (Mann-Whitney, p<0.01) after 5 hours of incubation (Fig 6B, left). We also investigated the percentage of somatic area covered by microglial processes over time, which has doubled at 0 minutes (fixed immediately after cutting) compared to the perfused controls (from 4% to 8%, p<0.05). This observation indicates extremely rapid microglia actions taking place during the ∼1-3 minutes between brain extraction and slice fixation. The massive initial increase was followed by a gradual decrease dropping back again to control values (purple indication; p<0.05) after 5 hours of incubation (Fig 6B, right). In line with this, microglia-synapse contact prevalence showed similar alterations after slice preparation in the case of glutamatergic synapses (Fig 6D, left), where average percentage of contacts increased by 46% between the perfused and 0-minute conditions (p<0.05) and gradually dropped back to control values. In the case of GABAergic synapses (Fig 6D, right), we did not observe significant increase immediately after slice preparation, however we measured a prominent decrease in contacts during the incubation process, with a 36% drop after 2 hours of incubation (p<0.01). Based on these observations, we concluded that microglia-neuron interaction sites underwent rapid and progressive changes, as somatic coverage and contact prevalence on glutamatergic synapses significantly increased immediately after slice preparation, followed by a gradual decrease over time.

**Figure 6.**
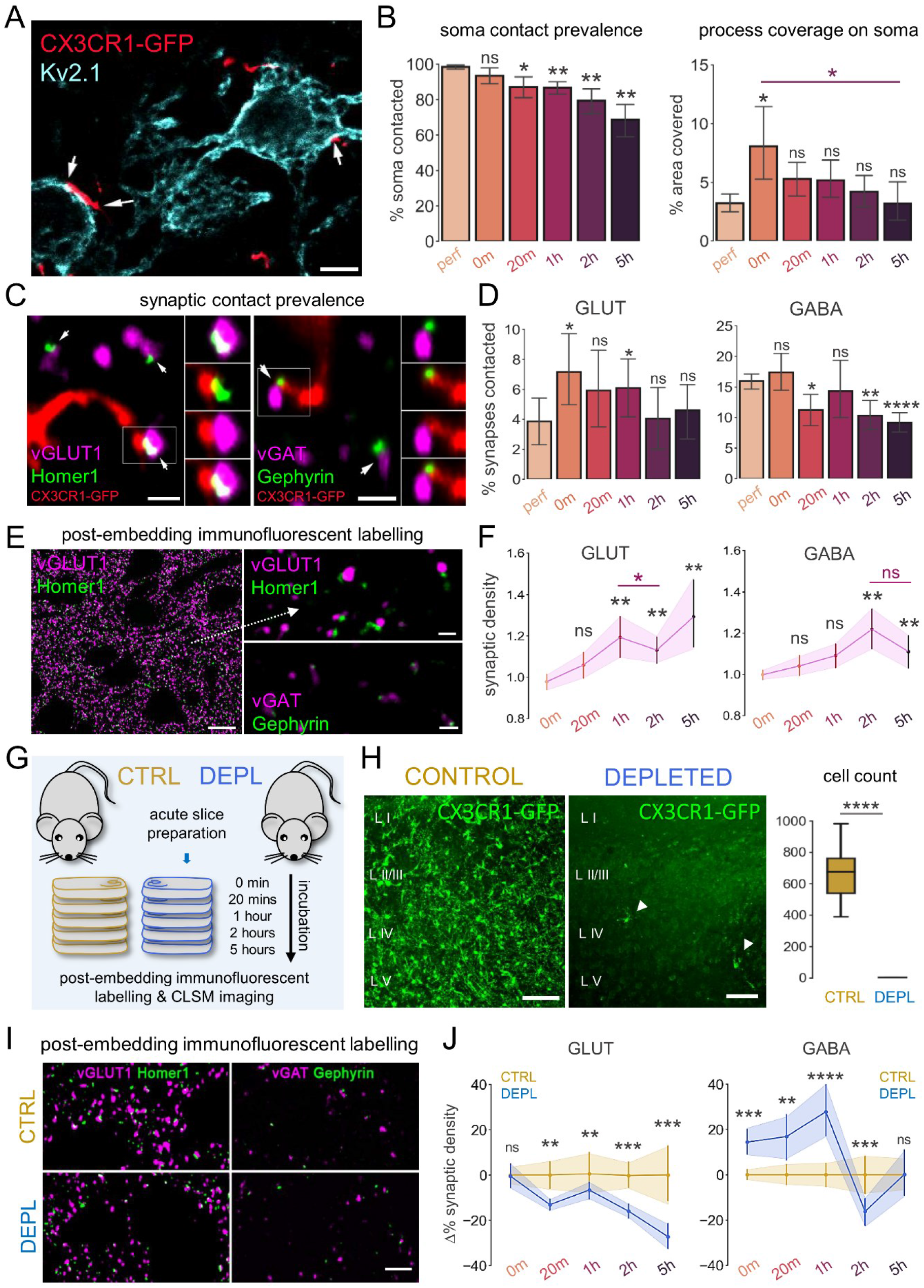
Microglial phenotype changes are accompanied by rapid alterations of microglia-neuron interactions and microglia-dependent synaptic sprouting. A. Representative section of a maximum intensity projection image created from confocal z-stacks used to quantify microglial contact prevalence and process coverage on neuronal soma. White arrows point to areas (overlap of microglia and Kv2.1 labelling) where microglial processes are likely to form contacts on neuronal soma (Scale bar: 5 µm). B. Quantification of contact prevalence (left) and coverage (right) of neuronal soma by microglial processes (left). N=3 animal, p.n.: ∼90 days; Mann-Whitney, ns: not significant, *: p<0.05, **: p<0.01, ***: p<0.001. Black statistical indication: perfused values vs. 0 min – 5 hours. Purple statistical indication: 0 min vs. 5 hours. C. Single image planes from confocal z-stacks used to quantify contact prevalence of microglial processes onto glutamatergic (left) or GABAergic (right) synapses. (Inserts show image channel pairs from the boxed area, from top to bottom: pre-and postsynaptic marker, microglia and postsynaptic marker, microglia, and presynaptic marker, merged.) Arrows show identified individual synapses contacted by microglial processes (bars: 1 µm). D. Quantification of microglial contact prevalence onto glutamatergic (left) or GABAergic (right) synapses. N=3 animal, p.n.: ∼90 days; Mann-Whitney, ns: not significant, *: p<0.05, **: p<0.01, ****: p<0.0001. E. Representative sections of maximum intensity projection images created from confocal z-stacks used to quantify glutamatergic and GABAergic synaptic densities. Images were created using the post-embedding labelling technique. A large area of glutamatergic synaptic labelling is shown (left, bar: 5 µm) and zoomed-in insets of glutamatergic (right, top; bar: 1 µm) and GABAergic synaptic labelling (right, bottom; bar: 1 µm). F. Quantification of synaptic density changes during the incubation process regarding glutamatergic (left) and GABAergic (right) synapses. N=6 animal, p.n.: ∼65 days; Mann-Whitney, ns: not significant, *: p<0.05, **: p<0.01, ***: p<0.001. G. Schematic representation of experiment. CX3CR1^+/GFP^ littermates were used to create a control (CTRL; N=3, p.n.: ∼65; brown) and microglia depleted (DEPL; N=3, p.n.: ∼65; blue) subgroup of animals. Slice preparations were obtained from both groups and immersion-fixed at different timepoints during recovery. H. Maximum intensity projection images showing microglial cells in acute slice preparations obtained from animals that either belong to control (left) or depleted (middle) group (bars: 100 µm). Quantification comparing the total number of microglial cells counted in control (CTRL, brown) or depleted (DEPL, blue) acute slices (right). N=6-6 animal, p.n.: ∼65 days; Mann-Whitney, **: p<0.01. I. Representative sections of maximum intensity projection images created from confocal z-stacks used to quantify and compare densities of glutamatergic (left) or GABAergic (right) synapses in control (top row) or microglia depleted (bottom row) acute slices (bar: 5 µm). J. Comparison of synaptic density changes during the incubation process regarding glutamatergic (left) and GABAergic (right) synapses between control and microglia depleted animals. The percentage of changes compared to the average values measured in control mice (brown, data also shown in panel F) are shown with changes calculated for microglia depleted (blue) acute slices. N=6 animal, p.n.: 65 days; Mann-Whitney, ns: not significant, **: p<0.01, ***: p<0.001, ****: p<0.0001.

Given the observed changes in microglial contact prevalence on synapses, we next examined how the density of synaptic elements changed during the incubation process. We used the quantitative post-embedding labelling method to precisely determine both glutamatergic and GABAergic synaptic density changes (Fig 6E) in acute slices. We found significant increases in both glutamatergic (20% increase compared to 0 min, p<0.01) and GABAergic (9% increase compared to 0 min, p<0.05) synaptic density after 1 hour of incubation (Fig 6F), similarly to previous reports (Bourne et al., 2007; Kirov et al., 1999; Trivino-Paredes et al., 2019). Interestingly, measuring synapse density at multiple timepoints during the incubation process revealed robust time-dependent changes concerning the proportions of excitatory and inhibitory synapses. While we observed similar, gradual increase in glutamatergic and GABAergic synaptic density towards the 1- and 2-hour timepoints, there was a 1-hour lag in the case of GABAergic synapses (Fig 6F, left vs. right), which peaked after 2 hours of incubation (20% increase compared to 0 min, p<0.001). Coincidentally, we observed a significant drop in glutamatergic synaptic density which occurred between 1- and 2 hours of incubation (Fig 6F left, purple statistical indication; 9% drop, p<0.05). We observed the highest density of glutamatergic synapses after 5 hours (28% increase compared to 0 min, p<0.01), whereas at this timepoint there was a tendency of decrease in GABAergic synaptic density (Fig 6F right; pink statistical indication; 10% drop, n.s., p=0.051). Based on these observations, we concluded that the neuronal network is likely to be reorganized by the slow and gradual build-up of excitatory synapse numbers, which is followed by the inhibitory synapses in the same manner. The inhibitory sprouting seemed to reach its maximum after 2 hours of incubation, while excitatory synapse numbers further increased towards the 5-hour timepoint.

Increasing evidence suggests that microglial cells are essential for the formation, pruning and maintenance of synapses both in the developing and in the adult CNS (Ikegami et al., 2019). We wanted to examine whether microglia can also actively contribute to the observed time-dependent structural and functional changes in synaptic density detected in acute slice preparations. To this end, we measured changes of synaptic densities in acute slice preparations in the absence of microglia (induced by elimination of microglia with PLX5622 for three weeks *in vivo* prior to slice preparation, Fig 6G-H). Depletion of microglia with PLX3373 or PLX5622 *in vivo* has previously been shown to induce no substantial changes in neuronal cell numbers and morphology, nor did it cause any deficits in cognition and behaviour, although dendritic spine numbers showed a slight increase (Elmore et al., 2014; Han et al., 2017; Strackeljan et al., 2021). To measure and compare synaptic density changes between control (CTRL) and depleted (DEPL) condition, we used the previously described post-embedding labelling method (Fig 6I). We found that time-dependent synaptic density changes after slice preparation were markedly influenced by microglial actions (Fig 6J). To our surprise, the absence of microglia abolished the gradual increase of glutamatergic synaptic density observed under control conditions and resulted in significantly lower synapse densities from 20 minutes after slice preparation and onward, reaching 27% lower synaptic density values in DEPL compared to CTRL after 5 hours, (p<0.001, Fig 6D, left). Even more prominent differences were observed regarding GABAergic synaptic densities (Fig 6J, right). The absence of microglia led to a marked initial increase of GABAergic synaptic density, peaking at 1 hour, followed by significant drop at 2 hours. This course fundamentally differs from that observed under control conditions, which peaks at 2 hours (19% higher in CTRL, p<0.001, Fig 6J, right). These results clearly indicate that microglia differentially control excitatory and inhibitory synaptic sprouting in acute slice preparations, initiating glutamatergic, while repressing GABAergic synapse formation, during the early stages (<1hour) following acute slicing.

### The absence of microglia or microglial P2Y12 receptors dysregulates neuronal network activity in acute slice preparations

It is well known that slices need a recovery period after cutting, to enable reliable and stable electrophysiological measurements. In line with this, we observed significant increase in SWR parameters after 2 hours in control slices (Supp. Fig 4), which paralleled marked, time-dependent changes in GABAergic and glutamatergic synapse densities.

Because the absence of microglia resulted in markedly lower glutamatergic and GABAergic synaptic densities after 2 hours of incubation (Fig 6J), we set out to examine whether synchronous events are also positively affected by the presence of microglia and compared features of spontaneously occurring SWR activity measured from control (CTRL), or microglia depleted (DEPL) acute slice preparations (Fig 7A-B). Simultaneous recordings from both conditions allowed precise quantification and comparison of the differences concerning the occurrence of SWR activity between the two conditions. We observed that 18 out of 36 slices (50%) presented detectable SWR activity in CTRL, whereas only 4 out of 36 slices (11%) from DEPL (Fig 7C). This striking difference confirms that microglial actions are necessary for the emergence of physiological-like network activity in *ex vivo* slices. Furthermore, our results showed significant differences between spontaneously occurring SWR activity recorded from CTRL or DEPL acute slice preparations, as we observed ∼3.5-fold decrease in SWR amplitude, ∼1.9-fold decrease in SWR rate and ∼2.1-fold decrease in ripple amplitude registered from DEPL when compared to CTRL slices (Fig 7D). These results indicate that microglia can effectively support the neuronal network in slice preparations during the incubation process, and can positively influence the occurrence, amplitude, and frequency of spontaneously occurring SWR activity.

**Figure 7.**
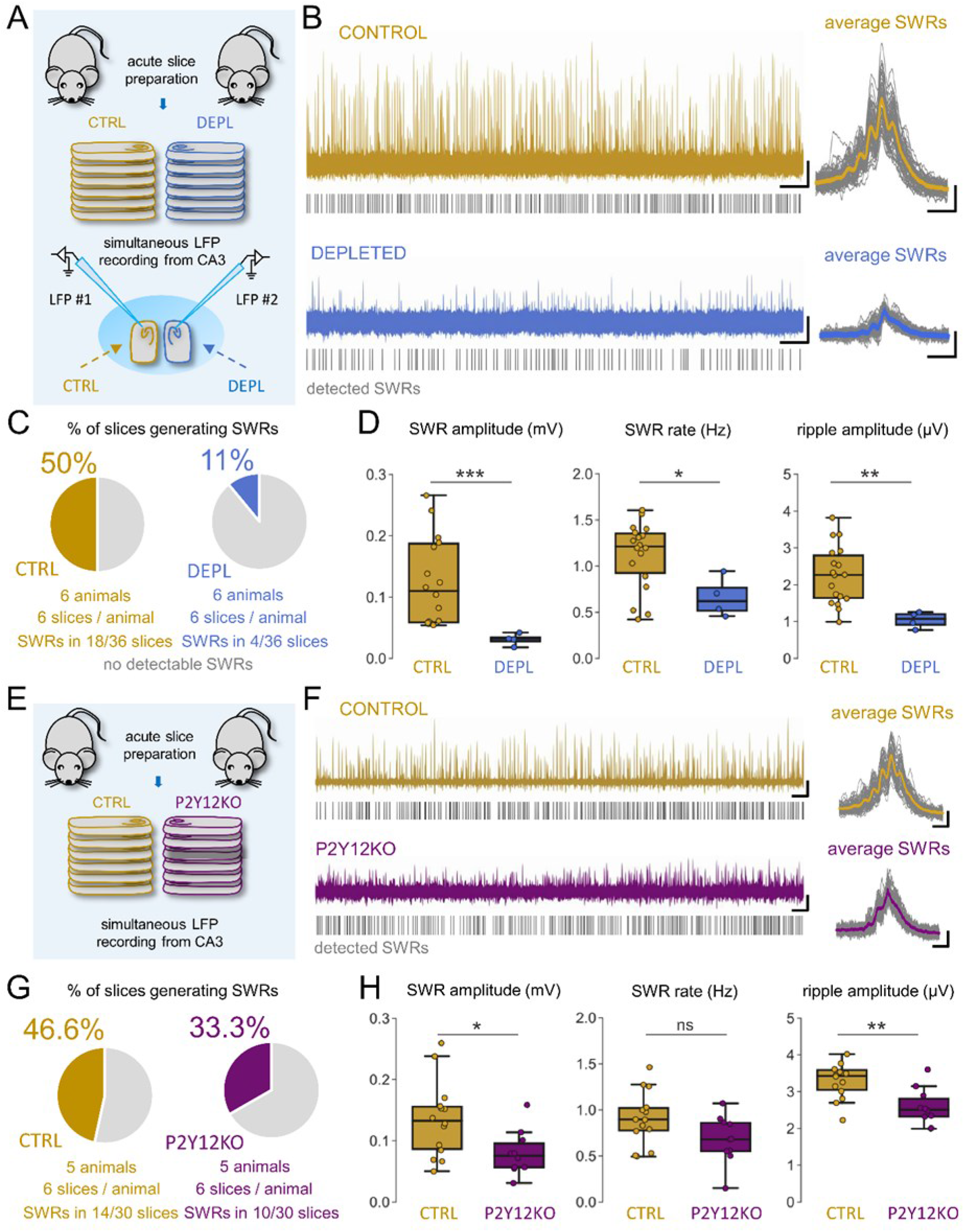
The absence of microglia or microglial P2Y12 receptors dysregulates neuronal network activity in acute slice preparations. A. Schematic representation of experiment. CX3CR1^+/GFP^ littermates were subjected to 3 weeks of either control or PLX3397 containing diet to create a control (CTRL; N=6, p.n.: ∼65; brown) and microglia depleted (DEPL; N=6, p.n.: ∼65; blue) subgroup of animals. Acute hippocampal slice preparations were obtained from both groups and placed into an interface-type incubation chamber for at least 1 hour of recovery. Subsequently, slices were transferred together in a pairwise manner into a recording chamber, to simultaneously measure sharp wave-ripple (SWR) activity via local field potential recordings (LFP) registered from the CA3 pyramidal layer of the hippocampus. B. Representative LFP recordings (left; bar: 30 s, 50 µV) and averages of spontaneous SWR events (right; #50 in total, bar: 100 ms, 25 µV) registered from control (top row, green) and microglia depleted (bottom row, blue) slices. Grey lines represent detected SWR events. C. Pie-charts representing SWR activity occurrence in slices measured from control group (green, 18/ 36 slices) versus microglia depleted (blue, 4/36 slices). D. Quantification of SWR amplitude (left), rate (middle) and Ripple amplitude (right) comparing events measured from control (CTRL, green) or depleted slices (DEPL, blue). Quantification showed a median value of SWR amplitude (in mV, 1st quartile, 3rd quartile) 0.109 (0.058, 0.191) in the case of CTRL and 0.032 (0.021, 0.04) in the case of DEPL conditions (Mann-Whitney test, p<0.001). Concerning SWR rate, we measured (in Hz) 1.21 (0.86, 1.37) in CTRL condition, and 0.62 (0.47, 0.88) in case of DEPL condition (p<0.05). We measured the Ripple amplitude to be (in µV, 1st quartile, 3rd quartile) 2.27 (1.59, 2.86) in the case of CTRL and 1.07 (0.82, 1.23) in the case of DEPL conditions (p<0.01). N=6 animal/group, p.n.: ∼65 days; Mann-Whitney test, ns: not significant, *: p<0.05, **: p<0.01, ***: p<0.001. E. Schematic representation of experiment. Control (CTRL; N=5, p.n.: ∼65; brown) and P2Y12 KO mice (DEPL; N=6, p.n.: ∼65; purple) were examined during the experiments, similarly as described in A. Acute hippocampal slice preparations were obtained from both groups and placed into an interface-type incubation chamber for at least 1 hour of recovery. Subsequently, slices were transferred together in a pairwise manner into a recording chamber, to simultaneously measure sharp wave-ripple (SWR) activity via local field potential recordings (LFP) registered from the CA3 pyramidal layer of the hippocampus. F. Representative LFP recordings (left; bar: 30 s, 50 µV) and averages of spontaneous SWR events (right; #50 in total, bar: 100 ms, 25 µV) registered from control (top row, brown) and P2Y12 KO (bottom row, purple) slices. Grey lines represent detected SWR events. G. Pie-charts representing SWR activity occurrence in slices measured from control group (brown, 14/ 30 slices) versus P2Y12 KO (purple, 10/30 slices). H. Quantification of SWR amplitude (left), rate (middle) and Ripple amplitude (right) comparing events measured from control or P2Y12 KO slices. Quantification showed a median value of SWR amplitude (in mV, 1st quartile, 3rd quartile) 0.133 (0.087, 0.155) in the case of CTRL and 0.075 (0.057, 0.096) in the case of P2Y12 KO (Mann-Whitney test, p<0.05). Concerning SWR rate, we measured (in Hz) 0.89 (0.77, 1.02) in CTRL, and 0.68 (0.55, 0.85) in case of KO (p=0.14). We measured the ripple amplitude to be (in µV) 3.42 (3.04, 3.58) in the case of CTRL and 2.51 (2.32, 2.81) in the case of KO (p<0.01). N=5 animal/group, p.n.: ∼65 days; Mann-Whitney test, ns: not significant, *: p<0.05, **: p<0.01.

**Supplementary Figure 4.**
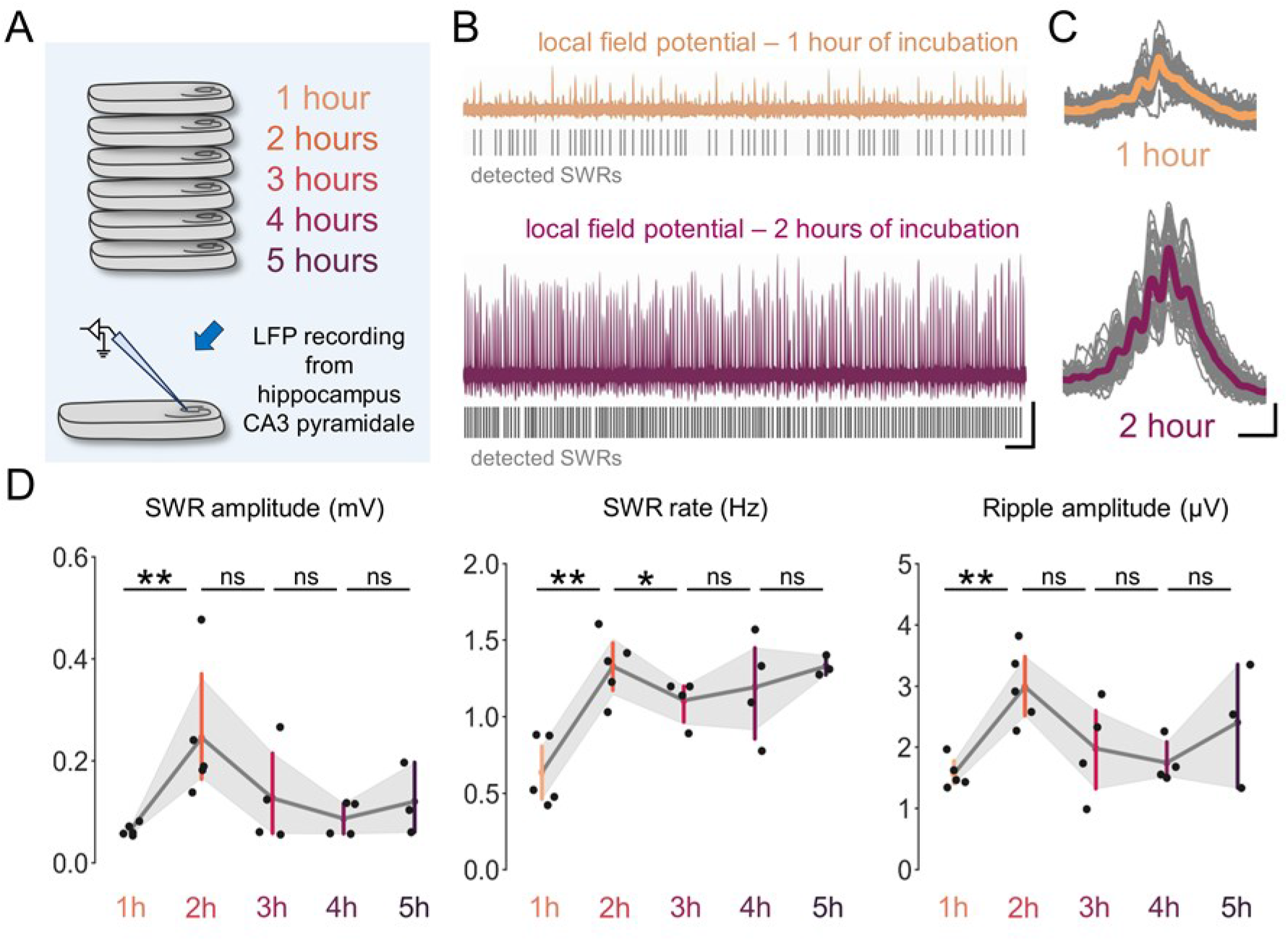
Temporal course of the emergence of Sharp Wave-Ripple activity after acute slice preparation. A. Schematic representation of the experiment. CX3CR1^+/GFP^ littermates (N=8; p.n.: ∼65 days) were used to create acute hippocampal slice preparations and placed into an interface-type incubation chamber for at least 1 hour of recovery time. Subsequently, slices were transferred at specific timepoints (hourly after 1-5 hours) into a recording chamber to measure sharp-wave ripple (SWR) activity via local field potential recordings (LFP) registered from the CA3 pyramidal layer of the hippocampus. B. Representative LFP recordings measured after 1 (yellow) or 2 hours (purple) of incubation. Grey lines represent detected SWR events (bars 10 s, 50 µV). C. Representative averaged traces of detected SWRs (#50 in total) measured after 1 hour (yellow: average, grey: individual events) or 2 hours (purple: average, grey: individual events) of incubation (bars 100 ms, 25 µV). D. Quantification of SWR amplitude (left), rate (right) and Ripple amplitude (right) comparing events measured after different time spent in the incubation chamber before recording. N=8 animal, p.n.: 65 days; Kruskal-Wallis test, ns: not significant, *: p<0.05, **: p<0.01.

To examine whether P2Y12 mediated actions are contributing to the observed effects, we repeated the same experiment by comparing slices from control and P2Y12 KO mice (Fig 7E-F). Similarly to microglia depleted mice, SWR occurrence was lower in the absence of P2Y12Rs (Fig 7G, 33%, 10 slices from 30 in total) when compared to control slices (46.6%, 14 slices out of 30). In this case, we also observed significant differences in SWR parameters, as SWR amplitude in average showed a ∼1.7-fold decrease (Fig 7H, left), while SWR ripple amplitude showed ∼1.3-fold decrease in P2Y12 KO animals (Fig 7H, right). SWR rate also decreased ∼1.3-fold compared to control slices, however due to a higher variance, this difference was not significant (Fig 7H, middle). Taken together, these results suggest that the positive influence of microglia on SWR activity observed in acute slice preparations is in part mediated by microglial P2Y12R action.

## Discussion

In this study, we provide a comprehensive assessment of microglial function in acute slice preparations in an experimentally relevant timeframe, using sample sets from multiple expert laboratories. We show that while microglia are undergoing marked time-dependent phenotype changes, they are instrumental to maintain *ex vivo* spontaneous activity of the neuronal network in a P2Y12R-dependent manner. We believe that these fundamental mechanisms of microglia-neuron interactions should be considered for studies on microglial and neuronal functions *ex vivo*, while also have important implications for the acute slice methodology and could facilitate improvements in modelling.

We first studied how the slice preparation procedure affected microglial cells in acute slices. While locomotor and process polarization behaviour of microglia in slice preparations have been previously suggested (Petersen & Dailey, 2004; Stence et al., 2001), population level changes in the distribution of cell bodies and processes along the full depth of slices have not been investigated in a time-dependent manner before. Because microglia are extremely sensitive to tissue disturbance, the acute slice preparation technique can ultimately be considered as a traumatic event assumed to influence microglial function, as observed in acute slices prepared via different methodologies (Kettenmann et al., 2011; Matyash et al., 2017; Petersen & Dailey, 2004; Stence et al., 2001). Of note, in comparison with other studies, microglia after 1 hour of incubation showed extremely similar morphological characteristics to cells measured at peri-infarct cortical areas in experimental stroke models (Heindl et al., 2018; Morrison & Filosa, 2013; Sadler et al., 2020; Singh et al., 2018). Moreover, the difference in the number of microglial process endings observed in a study comparing control and Alzheimer’s disease patients (Davies et al., 2017), closely resembles the changes that we observed in acute slices (∼50% drop) as early as after 20 minutes of incubation.

It has been previously suggested that gradually decreasing levels of oxygenation along the depth of the slice, as well as higher neuronal activity at the top surface could mediate injury-related signals (Huchzermeyer et al., 2013; Ivanov & Zilberter, 2011; Mulkey et al., 2001). Importantly, microglial phenotype changes were found to represent a generic response to slice preparation, as different slice preparation techniques carried out in independent laboratories consistently resulted in a gradually increasing density of reactive microglia both at slice surfaces and in deeper layers. While our observations reinforce general guidelines in electrophysiological investigations to avoid measurements taking place at the top 30-50 µm of slice preparations (Avignone et al., 2019), they provide novel mechanisms explaining the marked regional-and time-dependent heterogeneities observed in acute slices. Microglial recruitment to the site of injury and morphological transformation are known to be driven by extracellular ATP (which quickly hydrolyses to ADP) and P2Y12R actions (Davalos et al., 2005; Haynes et al., 2006; Nasu-Tada et al., 2005), which we assumed to take place upon slice preparation. To this end, we used a newly developed mouse line, where glutamatergic neurons express fluorescent ATP-sensor in the plasma membrane, enabling the visualisation of extracellular ATP-levels throughout the neuropil. While our results showing massive ATP-levels close to the cut slice surfaces were somewhat expected, we also reveal that the ATP-gradients are likely to drive microglial dislocation and morphological changes via P2Y12R throughout the slices, including deeper tissue locations. Initial ATP release effects could be further augmented by sustained ATP-flashes deep in the slices even hours after slice cutting. Such flashing ATP activity was also observed after acute brain injury *in vivo* (Chen et al., 2022). Important questions for future research will be how these local changes in extracellular ATP levels could influence microglial and neuronal physiology in *ex vivo* slice preparations.

P2Y12 receptors (P2Y12R) not only represent a core marker for microglia in the healthy brain (Bosco et al., 2018; Hickman et al., 2013; Peng et al., 2019), but downregulation of P2Y12R is associated with reactive phenotypes and microglial dysfunction both *in vivo* and *ex vivo* (Cserép et al., 2020; Gu et al., 2016; Haynes et al., 2006; Lin et al., 2021; Zrzavy et al., 2017). We show with a fully quantitative post-embedding method (Holderith et al., 2020) that rapid downregulation of P2Y12R occurs most extensively at the processes of microglia already after 1 hour of incubation proportionally with marked morphological changes. Resting membrane potential of microglia has also been shown to correlate with morphology, as modified activity of tonically active K+ channels responsible for maintaining resting membrane potential (knocking out THIK-1 or locally increasing extracellular [K+]) can decrease ramification (Madry, Arancibia-Cárcamo, et al., 2018; Madry, Kyrargyri, et al., 2018). It has been proposed that transitioning into a more reactive phenotype could partly reflect decreased expression of THIK-1 (Madry, Kyrargyri, et al., 2018), as activation via LPS treatment resulted in a significantly downregulated expression of THIK-1 mRNA (Holtman et al., 2015). Importantly, our results indicate that gradual depolarization, rapid P2Y12R downregulation and transitioning into a reactive morphology are co-occurring events in acute slice preparations solely at “baseline” conditions, which have important implications for models using acute slice preparations.

Changes in microglial distribution, morphology, P2Y12R-expression and local ATP fluxes are expected to markedly influence cell-cell interactions, given that P2Y12R signalling is required for normal microglial interactions with neuronal somata and synapses under different conditions (Cserép et al., 2020; Lin et al., 2021; Miyamoto et al., 2016; Sipe et al., 2016). We showed that while the prevalence of contacts on neuronal soma (somatic junctions) slowly and gradually decreases, the percentage of somatic area covered by microglial processes undergoes a more than two-fold increase immediately after slice preparation similarly to that observed *in vivo* after acute ischemia (Cserép et al., 2020). These changes paralleled increased numbers of contacts on glutamatergic synapses, while contacts on GABAergic synapses were gradually decreasing. Studies have previously demonstrated that acute slice preparation induces synaptic sprouting (Bourne et al., 2007; Kirov et al., 1999; Trivino-Paredes et al., 2019), but the mechanisms of these changes have not been revealed. Microglia are well-known regulators of synaptic density *in vivo* (Akiyoshi et al., 2018; Miyamoto et al., 2013, 2016; Schafer et al., 2013; Wake et al., 2009; Y. Wu et al., 2015). Of note, we found that the absence of microglia profoundly influenced the course of synaptic sprouting. While microglial contacts can facilitate spine formation of functional synapses both during development and in adult stages (Miyamoto et al., 2013, 2016; Wake et al., 2009; Weinhard et al., 2018; Y. Wu et al., 2015) the extent and speed of microglia-mediated effects in *ex vivo* brain slices was surprising. Thus, further studies will be required to reveal the molecular mechanisms underlying these effects, potentially making the acute slice maturation process a valuable model for studying microglia-synapse interactions.

To our knowledge, we show for the first time that the presence of functional microglia can actively influence SWR activity in hippocampal acute slice preparations, which was largely mediated by purinergic mechanisms. Thus, the presence of microglia and functional P2Y12R signalling are required for the emergence of physiological-like network activity in *ex vivo* slices suggesting that interventions aiming to inhibit microglial actions could render acute brain slices less capable of producing normal patterns of neuronal network activity seen *in vivo*. Previously it has been demonstrated that hippocampal field excitatory postsynaptic potentials (fEPSPs) recover in correlation with synaptic changes in *ex vivo* slices (Kirov et al., 1999). Our results show that SWR activity shows a huge increase in amplitude, rate and frequency when correlated to the measured synaptic density changes. Since the generation of SWRs in these conditions are both reliant on a sufficiently large tonic excitatory activity and the action of reciprocally connected parvalbumin positive basket cells (Schlingloff et al., 2014), these results indicate that a delicate balance needs to be maintained between excitation and inhibition during the course of sprouting which allows SWRs to emerge. Importantly, our results showed that microglia can facilitate glutamatergic and repress GABAergic synaptic formation to maintain a balanced reorganization of the network after the loss of synapses due to the slice preparation procedure. Supporting our data, partial depletion of microglia promoted asynchronous activity without overall change in frequency (Akiyoshi et al., 2018), while microglia depletion in the hippocampus decreased spontaneous and evoked glutamatergic activity (Basilico et al., 2022). In contrast, *in vivo* imaging experiments showed increased excitatory and inhibitory neuronal activity in the cortex after microglia depletion, which results are in line with *ex vivo* circuit mapping data (Császár et al., 2022; Liu et al., 2021, 2022). Thus, it is likely that in addition to the synaptic density changes, the altered distribution, morphology, and subsequent changes in microglia-neuron interactions are also key players in maintaining the integrity of neuronal networks in *ex vivo* acute slices, enabling complex, *in vivo*-like network activity patterns to emerge.

Acute slice preparation is a well-established experimental approach, which is extremely useful to study the physiology of neurons and other cells from individual synapses to complex neuronal networks. However, precautions are necessary during the interpretation of results, as certain changes in these conditions may reflect different physiological or pathological states concerning both neuronal and microglial functions. In fact, we show that microglia in acute slices can present a rapid transition into a more reactive phenotype and can actively influence the neuronal network through interactions via altered microglia-neuron interactions and purinergic signalling. Since microglial morphology strongly reflects the state of the tissue, we suggest that monitoring these changes should always be considered, as it could help in the contextualization of results concerning both the microglial and neuronal populations. Moreover, an extensive examination and comparison of microglial phenotype state changes due to different slice preparation methods could facilitate more refined and consistent experimental models and paradigms to be established. In addition, our results also emphasize the importance of interactions between microglia and complex neuronal networks, which may be further emphasized by the sensitivity of microglia to a broad range of changes in their micro-and macroenvironment. While the observed changes in microglia in acute slices may not represent an undisturbed physiological state, the acute slice model also emerges as an instrumental tool to test and understand the different factors, which contribute to reactive microglial phenotypes, with broad implications for diseases of the CNS that are influenced by alterations of microglial function.

## Materials & Methods

### Animals

In all experiments, CX3CR1^+/GFP^ or C57Bl/6J mice littermates of both sexes were used. Mice were kept in the vivarium on a 12-hour light/dark cycle and provided with food and water ad libitum. The animals were housed two or three per cage. All experiments were approved by the Ethical Committee for Animal Research at the Institute of Experimental Medicine, Hungarian Academy of Sciences, and conformed to Hungarian (1998/XXVIII Law on Animal Welfare) and European Communities Council Directive recommendations for the care and use of laboratory animals (2010/63/EU) (license number PE/EA/2552-6/2016; PE/EA/254-7/2019).

### Slice preparation, incubation and fixation

In order to minimize bacterial contamination, all the tools and containers used for slice preparation and incubation were routinely cleaned before and after experiments with 70% ethanol and were rinsed extensively with distilled water. For acute slice preparation, mice were decapitated under deep isoflurane anesthesia. The brain was removed and placed into an ice-cold cutting solution, which had been bubbled with *95% O2–5% CO2* (carbogen gas) for at least 30 min before use. The cutting solution contained the following (in mM): *205 sucrose, 2.5 KCl, 26 NaHCO3, 0.5 CaCl2, 5 MgCl2, 1.25 NaH2PO4, 10 glucose*, saturated with *95% O2–5% CO2*. Horizontal hippocampal slices of 300 µm or 450 µm (in case of LFP recordings) thickness were cut using a Vibratome (Leica VT1000S). The process of slice preparation from termination till the first slice to be immersion-fixed took ∼5-10 minutes.

After acute slice preparation, slices were placed into an interface-type holding chamber for recovery. In an interface-type chamber, slices are laid onto a mesh just slightly submerged into the artificial cerebrospinal fluid (ACSF), therefore the oxygenation of the tissue is mainly realized by the direct exposure to humidified oxygen-rich air above the slices. This chamber contained standard ACSF at 35°C that gradually cooled down to room temperature. The ACSF solution contained the following (in mM): *126 NaCl, 2.5 KCl, 26 NaHCO3, 2 CaCl2, 2 MgCl2, 1.25 NaH2PO4, 10 glucose*, saturated with *95% O2–5% CO2*. Immediately after slice preparation/given timeframes of incubation/after recordings, slices were immersion-fixed for 1 hour with 4% PFA solution. In the case of PSB treated acute slices, both cutting and standard ACSF solutions used to prepare the slices contained 10 µM PSB- 0739 (Sigma-Aldrich).

For perfusion-fixed slices, mice were anesthetized and transcardially perfused with 0.9% NaCl solution for 1 minute, followed by 4% PFA in 0.1 M phosphate buffer (PB) for 40 minutes, followed by 0.1 M PB for 10 minutes to wash the fixative out. Blocks containing the somatosensory cortex and ventral hippocampus were dissected, and horizontal sections were prepared on a vibratome (VT1200S, Leica, Germany) at 50 μm thickness for immunofluorescent histological and 100 μm thickness for the automated morphological analysis.

#### Lab#2 slice preparation protocol

Mice were anesthetized with isoflurane and decapitated according to a protocol approved by the UCLA Chancellor’s Animal Research Committee. After decapitation the head was immersed in ice-cold ACSF solution (see below) and put in the −80°C freezer for 1 minute. The brain was then removed from the skull and coronal 350 μm thick slices were cut from a range of AP (from bregma): - 0.5 to −3 mm on a Leica VT1000S vibratome in ice-cold N-Methyl-D-Glutamine (NMDG)-based HEPES-buffered solution, containing (in mM): 135 NMDG, 10 D-glucose, 4 MgCl2, 0.5 CaCl2, 1 KCl, 1.2 KH2PO4, 20 HEPES, 27 sucrose (bubbled with 100% O2, pH 7.4, 290-300 mOsm/L). Then, slices were incubated at 34°C in a reduced sodium + sucrose artificial CSF (ACSF), containing (in mM): NaCl 85, D-glucose 25, sucrose 55, KCl 2.5, NaH2PO4 1.25, CaCl2 0.5, MgCl2 4, NaHCO3 26, pH 7.3-7.4 when bubbled with 95% O2, 5% CO2 before fixation. Each slice was placed with the same orientation into an incubation chamber, and the slice surface was identified as the side facing towards the top. The incubation period for a series of slices from the same animal was designed as: 0 minutes (this slice was also placed into the incubation chamber for 3-5 seconds), 20 minutes, 1 hour, 2 hours and 5 hours. After a certain time of incubation, the corresponding slice was carefully transferred from the incubation chamber to a 24 well plate pre-filled with 4% freshly depolimerized paraformaldehyde in 0.1 M phosphate buffer (PFA) for fixation. Each slice spent exactly 1 hour in fixative before transferring into the “wash” plate and washed extensively with 0.1 PBS by exchanging the PBS in the well 3 times first, then placing the washing plate on a shaker and washing in PBS for 3×10 minutes. Slices were then individually inserted into 15 ml plastic test-tubes sandwiched between tissue paper to avoid shaking during transportation. The test-tubes were placed in a Styrofoam container and shipped to the IEM.

#### Lab#3 slice preparation protocol

C57Bl/6J mice at the age of 4 weeks were decapitated under deep isoflurane anesthesia. The brain was quickly removed and immersed in ice-cold sucrose-containing artificial cerebrospinal fluid, saturated with 95% O2- 5% CO2 (sucrose-containing ACSF; in mM: NaCl 60, sucrose 100, KCl 2.5, NaH2PO4 1.25, NaHCO3 26, CaCl2 1, MgCl2 5, D-glucose 20; pH 7.4, 310 mOsmol). Coronal hippocampal slices (300 µm thick) were prepared with a vibratome (VT1200S, Leica). The slices were incubated at 35℃ in the same solution for 20 min. The 0-minute slice was placed into the incubation chamber for a few seconds (3-5 seconds) before fixation with 4% PFA. Then, 20 minutes slices were fixed with 4% PFA before being transferred to second chamber filled with ACSF saturated with 95% O2- 5% CO2 at room temperature (ACSF; in mM: NaCl 125, KCl 3.5, NaH2PO4 120, NaHCO3 26, CaCl2 2, MgCl2 5, D-glucose 15; pH 7.4, 310 mOsmol). After that, 1 hour, 2 hours and 5 hours slices were fixed with 4% PFA. Each slice spent 1 hour for fixation, and then transferred into wash plates containing 0.1 M phosphate buffer (PB) and washed with PB 3 times for 10 minutes. All slices were stored and transported submerged under 0.1 M PB, supplemented with 0.05 % sodium-azide.

### Cross section of slice preparations and quantification of translocation

300 µm thick acute slices were immersion fixed immediately after slicing (0 minute) or after 20 minutes, 1 hour, 2 hours or 5 hours spent in an interface-type incubation chamber (Fig 1A). Fixed slices were washed in 0.1M PB, flat embedded in 2% agarose blocks, rotated 90 degrees, and resliced on a vibratome (VT1200S, Leica, Germany) at 50 μm thickness (Fig 1B). The sections were mounted on glass slides, and coverslipped with Aqua-Poly/Mount (Polysciences). Intrinsic (CX3CR1^+/GFP^) immunofluorescence was analyzed using a Nikon Eclipse Ti-E inverted microscope (Nikon Instruments Europe B.V., Amsterdam, The Netherlands), with a CFI Plan Apochromat VC 20X DIC N2 objective (numerical aperture: 075) and an A1R laser confocal system. We used 488 nm excitation laser (CVI Melles Griot), and image stacks (resolution: 0.62 µm/px) were taken with NIS-Elements AR. Maximal intensity projections of stacks containing the whole section thickness were saved in tiff format, cell bodies were masked with Fiji “Analyze particles” plugin. Cell-body masks were used to count cells, and these masks were subtracted from the original tiff files to get images containing microglial processes only. As a validation for volume-related quantifications in acute slices, the average thickness for each preparation across the incubation procedure was measured (mean±SD: 262±7 µm), which showed no substantial differences between groups (Kruskal-Wallis test, p>0.05). For quantification, a measuring grid was placed onto the entire thickness of cross-sections (Fig 1 C), which divided the thickness into 7 equal zones. Cell body numbers were counted within these grids, and microglial process volume was assessed by measuring fluorescent integrated density within the grids with Fiji software.

Cell-body translocation calculation (Fig 1F) was performed for assessing microglial movement towards the top surface, and for away from the middle Z-depth of the slices. The coordinates for cell bodies were registered in Fiji software, the distance of the cell bodies from the bottom surface or the (middle Z-depth) were measured at 0 min, and at 5 hours (the number of measured cells was identical at the two time-points). The distances were sorted in growing rows for both timepoints, and the 0-minute values subtracted from the 5-hour values, thus we could calculate the minimal values cells had to travel in order to reach the final distribution pattern (at 5 hours) starting from the 0 minutes distribution. For the process area coverage measurement (Fig 1I) the images with cell bodies masked out were used. Images from slices fixed at 0 minutes and 5 hours were binarized in Fiji, and the percentage of covered area measured.

### Time-lapse imaging

Acute brain slices (300µm thick) were prepared from 80-day old CX3CR1^+/GFP^ mice as described above. Z-stack images (1µm step size) were acquired using a Nikon C2 laser scanning confocal microscope equipped with a 20x CFI Plan Apo VC (NA=0.75 WD=1.00mm FOV=1290.4mm) objective at 488nm, under continuous perfusion with ACSF (3 ml/min perfusion rate). The image acquisition started 1h after slice cutting. Image stacks were taken every 20 min for 6 hours. Video editing was performed using NIS Elements 5.00 and ImageJ 1.53f51.

### Automated morphological analysis of microglial cells

300 µm thick acute slices were immersion fixed for 1 hour immediately after slicing (0 minute) or after 20 minutes, 1 hour, 2 hours or 5 hours spent in an interface-type incubation chamber (Fig 1A). Fixed slices were washed in 0.1M PB, flat embedded in 2% agarose blocks, and re-sectioned on a vibratome (VT1200S, Leica, Germany) at 100 μm thickness (Fig 1B). Sections selected from the middle region of incubated slices were immunostained with antibodies, and DAPI (for primary and secondary antibodies used in this study, please see Table 1.). Preparations were kept in free-floating state until imaging to minimize deformation of tissue due to the mounting process. Imaging was carried out in 0.1M PB, using a Nikon Eclipse Ti-E inverted microscope (Nikon Instruments Europe B.V., Amsterdam, The Netherlands), with a CFI Plan Apochromat VC 60X water immersion objective (numerical aperture: 1.2) and an A1R laser confocal system. Volumes were recorded with 0.2 µm/pixel resolution and a Z-step of 0.4µm. For 3-dimensional morphological analysis of microglial cells, the open-source MATLAB-based Microglia Morphology Quantification Tool was used (available at https://github.com/isdneuroimaging/mmqt). This method uses microglia and cell nuclei labeling to identify microglial cells. Briefly, 59 possible parameters describing microglial morphology are determined through the following automated steps: identification of microglia (nucleus, soma, branches) and background, creation of 3D skeletons, watershed segmentation and segregation of individual cells (Heindl et al., 2018).

**Table 1.** List of antibodies used in the study.

| **Primary antibodies** | **host** | **source** | **catalog nr.** | **RRID** |
| --- | --- | --- | --- | --- |
| Gephyrin | Mouse | Synaptic Systems | 147 021 | AB_2232546 |
| GFP | Chicken | Invitrogen | A10262 | AB_2534023 |
| Homer1 | Rabbit | Synaptic Systems | 160 003 | AB_887730 |
| IBA1 | Guinea pig | Synaptic Systems | 234004 | AB_2493179 |
| Kv2.1 | Mouse | NeuroMab | 75-014 | AB_10673392 |
| P2Y12R | Rabbit | Anaspec | AS-55043A | AB_2298886 |
| TMEM119 | Chicken | Synaptic Systems | 400 006 | AB_2744643 |
| vGAT | Guinea pig | Synaptic Systems | 131 004 | AB_887873 |
| VGLUT1 | Guinea pig | Synaptic Systems | 135 304 | AB_887878 |
| **Secondary antibodies** | **host** | **source** | **catalog nr.** | **RRID** |
| Alexa 488 Streptavidin | - | - | S-11223 | - |
| Alexa 488 anti-chicken | donkey | Jackson ImmunoResearch Labs | 703-546-155 | AB_2340376 |
| Alexa 488 anti-guinea-pig | donkey | Jackson ImmunoResearch Labs | 706-546-148 | AB_2340473 |
| Alexa 488 anti-guinea-pig | donkey | Jackson ImmunoResearch Labs | 706-546-148 | AB_2340473 |
| Alexa 488 anti-mouse | donkey | Thermo Fisher Scientific | A-21202 | AB_141607 |
| Alexa 488 anti-rabbit | donkey | Jackson | 711-546-152 | AB_2340619 |
| Alexa 594 anti-guinea-pig | goat | LifeTech | A11076 | AB_141930 |
| Alexa 594 anti-mouse | donkey | Invitrogen | A-21203 | AB_141633 |
| Alexa 594 anti-rabbit | donkey | LifeTech | A21207 | AB_141637 |
| Alexa 647 anti-chicken | donkey | Jackson ImmunoResearch Labs | 703-606-155 | AB_2340380 |
| Alexa 647 anti-rabbit | donkey | Jackson ImmunoResearch Labs | 711-605-152 | AB_2492288 |
| Alexa 647 anti-guinea-pig | donkey | Jackson ImmunoResearch Labs | 706-606-148 | AB_2340477 |
| Biotinylated anti-chicken | goat | Vector Laboratories | BA-9010 | AB_2336114 |

### Pre-embedding immunofluorescent labelling and analysis of CLSM data

Before the immunofluorescent labelling, the 50 µm thick sections were washed in PB and Tris-buffered saline (TBS). Thorough washing was followed by blocking for 1 hour in 1% human serum albumin (HSA; Sigma-Aldrich) and 0.03-0.1% Triton X-100 dissolved in TBS. After this, sections were incubated in mixtures of primary antibodies, diluted in TBS overnight at room temperature. After incubation, sections were washed in TBS and were incubated overnight at 4 °C in the mixture of secondary antibodies, all diluted in TBS. Secondary antibody incubation was followed by washes in TBS, PB, the sections were mounted on glass slides, and coverslipped with Aqua-Poly/Mount (Polysciences). Immunofluorescence was analyzed using a Nikon Eclipse Ti-E inverted microscope (Nikon Instruments Europe B.V., Amsterdam, The Netherlands), with a CFI Plan Apochromat VC 60X oil immersion objective (numerical aperture: 1.4) and an A1R laser confocal system. We used 405, 488, 561 and 647 nm lasers (CVI Melles Griot), and scanning was done in line serial mode, pixel size was 50×50 nm. Image stacks were taken with NIS-Elements AR. For primary and secondary antibodies used in this study, please see Table 1. Quantitative analysis of each dataset was performed by at least two observers, who were blinded to the origin of the samples, the experiments, and did not know of each other’s results.

For the analysis of somatic contact prevalence, confocal stacks with double immunofluorescent labeling (cell type-marker and microglia) were acquired from at least three different regions of mouse cortex. All labeled and identified cells were counted when the whole cell body was located within the Z-stack. Given somata were considered to be contacted by microglia, when a microglial process clearly touched it (i.e., there was no space between neuronal soma and microglial process) on at least 0.5 µm long segment.

Microglial process coverage was measured on CLSM Z-stacks acquired with a step size of 300 nm. On single-channel images, Kv2.1-positive cells were selected randomly, the cell bodies of which were fully included in the captured volume. The surface of these cells was calculated by measuring the circumference of the soma on every section multiplied by section thickness. The surface of microglial process contacts was measured likewise.

For the analysis of synaptic contact prevalence, confocal stacks with triple immunofluorescent labeling (pre-and postsynaptic markers and microglia) were analyzed using an unbiased, semi-automatic method. First, the two channels representing the pre-and postsynaptic markers were exported from a single image plane. The threshold for channels were set automatically in FIJI, the „fill in holes” and „erode” binary processes were applied. After automatic particle tracking, synapses were identified where presynaptic puncta touched postsynaptic ones. From these identified points we selected a subset in a systematic random manner. After this, the corresponding synapses were found again in the original Z-stacks. A synapse was considered to be contacted by microglia, when a microglial process was closer than 200 nm (4 pixels on the images).

### Post-embedding immunofluorescent labelling and quantitative analysis

The technique described by Holderith et al. (Holderith et al., 2021.) was used with slight modifications. 300 µm thick acute slices were cut from CX3CR1^+/GFP^ mouse line and then immersion fixed immediately after slicing (0 minute) or after 20 minutes, 1 hour, 2 hours or 5 hours spent in an interface-type incubation chamber (Fig 1 B). Fixed slices were washed in 0.1M PB and 0.1M Maleate Buffer (MB, pH: 6.0). Then slices were treated with 1% uranyl-acetate diluted in 0.1M MB for 40 minutes in dark. This was followed by several washes in 0.1M PB, then slices were dehydrated in ascending alcohol series, acetonitrile and finally embedded in Durcupan (Fluca). Each block contained all slices from a respective time series of one animal. Ultrathin sections were cut using a Leica UC7 ultramicrotome at 200 nm thickness and collected onto Superfrost Ultra plus slides and left on a hotplate at 80°C for 30 minutes then in oven at 80°C overnight (Fig 3 C). Sections were encircled with silicon polymer (Body Double standard kit, Smooth-On, Inc.) to keep incubating solutions on the slides. The resin was etched with saturated Na-ethanolate for 5 minutes at room temperature. Then sections were rinsed three times with absolute ethanol, followed by 70% ethanol and then DW. Retrieval of the proteins were carried out in 0.02M Tris Base (pH = 9) containing 0.5% sodium dodecyl sulfate (SDS) at 80°C for 80 min. After several washes in TBS containing 0.1% Triton X-100 (TBST, pH = 7.6), sections were blocked in TBST containing 6% BlottoA (Santa Cruz Biotechnology), 10% normal goat serum (NGS, Vector Laboratories) and 1% BSA (Sigma) for 1 hour then incubated in the primary Abs diluted in blocking solution at room temperature overnight with gentle agitation. After several washes in TBST the secondary Abs were applied in TBST containing 25% of blocking solution for 3 hours. After several washes in TBST, slides were rinsed in DW then sections were mounted in Slowfade Diamond (Invitrogen) and coverslipped. Immunofluorescence was analyzed using a Nikon Eclipse Ti-E inverted microscope (Nikon Instruments Europe B.V., Amsterdam, The Netherlands), with a CFI Plan Apochromat VC 60X oil immersion objective (numerical aperture: 1.4) and an A1R laser confocal system. We used 488 and 647 nm lasers (CVI Melles Griot), and scanning was done in line serial mode, pixel size was 50×50 nm. Image stacks were taken with NIS-Elements AR. For primary and secondary antibodies used in this study, please see Table 1. Quantitative analysis of each dataset was performed by at least two observers, who were blinded to the origin of the samples, the experiments, and did not know of each other’s results.

For the quantitative assessment of P2Y12R expression, single high-resolution CLSM image planes were used. Microglial cell bodies, thick (average diameter greater than 1 µm) and thin (average diameter less than 1 µm) processes were identified based on TMEM119 and CX3CR1^+/GFP^ staining. Once the respective outlines of these profiles have been delineated, these outlines have been extended both in the intra-and the extracellular direction with 250-250 nm, yielding a 500 nm wide ribbon-shaped ROI. The integrated fluorescent density of P2Y12R-labeling was measured and divided by the lengths of the respective ROIs, which gave us the P2Y12R fluorescent intensity values applied to unit membrane lengths for each profile.

For the synapse density measurements, we used double stainings for pre-and postsynaptic markers (vGluT1 with Homer1 for glutamatergic, and vGAT with Gephyrin for GABAergic synapses). We could validate the specificity and sensitivity of these stainings based on the near-perfect match between pre and postsynaptic markers, thus we continued to measure the presynaptic signals. ROIs were randomly chosen within the neuropil, avoiding cell bodies. The integrated fluorescent densities were measured within these ROIs for vGluT1 and vGAT channels. Measurements were performed with the Fiji software package.

### Ex vivo 2-photon imaging of extracellular ATP

For extracellular expression of the ATP sensor GRABATP in neurons, Vglut1/cre/Gt(Rosa26)Sor[cre/cre] mice(ref) were crossed with LSL_GRABATP_P2A_jRGECO1a (TgTm) - C57Bl/6J [flox/flox] mice made by Biocytogen (China). Mice were used in the *ex vivo* 2- photon imaging experiments at the age of 67-72 days. Acute slices were obtained as in previous experiments (Methods/Slice preparation and incubation). Slices including the hippocampus and cortical regions above were used for two photon imaging (5 slices/brain). The slices were split and from each plane one half was used for imaging within the CA3 area in the hippocampus and the other half for imaging within layer two in the cortex, both in a 256*128 um window. The time between slice cutting and imaging was minimized to 8-13 min for the very first slices. For each slice, 10 min imaging at 0.5 frame/sec rate was followed by Z-stack acquisition (3 um step size; taking ∼3-4 min, see experimental setup on Fig 5A). Two photon imaging was accomplished by a Nikon Eclipse FN1 upright microscope, equipped with a Nikon A1R MP+ multiphoton system using a Chameleon Vision II Ti: Sapphire tunable (680-1080 nm) laser and a 25x water dipping objective lens [CFI75 Apo LWD NA=1.1 WD=2 mm]. The excitation wavelength was set to 920 nm for detecting the signal and 550/88 nm emission filter and a GaAsP NDD PMTs detector was used. Time lapse data were analyzed using the NIS-Elements (version 5.42.02) and ImageJ (version 1.53t) software. Mean fluorescent intensity (MFI) decreases (Fig 5B-C) were measured in areas where no ATP flash activity (focal ATP surges) was observed [ROI size: 12,5*12,5 um; 4 ROI/slice]. Z-stack images were used to determine the fluorescent signal decrease in time and depth (Fig 5D-E, Suppl Fig 2C), by averaging MFI data from 3 regions [ROI size: 25*50um] from each slice. ATP flashes were identified on each slice manually, encircling the individual flashes when the signal intensity was at maximum. Then, flash size, mean fluorescent intensity, peak duration, rise time and decay time data were collected and analyzed. Details of statistical analysis are provided in Fig 5 legend and Suppl Fig 3.

### Selective depletion of microglia

CX3CR1^+/GFP^ or C57B1/6J littermates were subjected to 3 weeks of either control or PLX3397 containing diet to create control and microglia depleted subgroup of animals, respectively. Extra slices were gathered from each animal by the re-slicing of immersion-fixed or perfusion-fixed acute slice preparations (50 µm). Success of depletion was monitored by creating z-stack images of the native GFP signal of microglia with confocal laser-scanning microscopy and verified by comparing the total number of microglia counted (via Fiji counting tool) at the same cortical and hippocampal locations, and through the whole depth of the slice preparations.

### Patch clamp recordings

Generally accepted guidelines were followed for patching microglial cells (Avignone et al., 2019). After incubation for given timeframes (as specified for each experiment in the Results section), slices were transferred individually into a submerged-type recording chamber with a superfusion system allowing constantly bubbled (95% O2–5% CO2) ACSF to flow at a rate of 3-3.5 ml/min. The ACSF was adjusted to 300-305 mOsm and was constantly saturated with 95% O2–5% CO2 during measurements. All measurements were carried out at 33 –34°C, temperature of ACSF solution was maintained by a dual flow heater (Supertech Instruments). The pipette solution contained (in mM): 120 KCl, 1 CaCl2, 2 MgCl2, 10 HEPES, and 11 EGTA, pH: 7.3, 280-300 mOsm. Pipette resistances were 3-6 MΩ when filled with pipette solution. Visualization of slices and selection of cells (guided by native GFP signal) was done under an upright microscope (BX61WI; Olympus, Tokyo, Japan equipped with infrared-differential interference contrast optics and a UV lamp). Only cells located deeper than ∼50 µm measured from the slice surface were targeted. All cells were initially in voltage-clamp mode and held at −40 mV holding potential during the formation of the gigaseal. Series resistance was constantly monitored after the whole-cell configuration was established, and individual recordings taken for analysis showed stability in series resistance between a 5% margin during the whole recording. After whole-cell configuration was established, resting membrane potential values were measured by changing the recording configuration to current-clamp mode at 0 pA for a short period of time (10-15 seconds) and evaluated from the recorded signal via averaging a 5 second period. Thereafter, responses to a pulse-train of current steps (−2 pA to −10 pA with 2 pA increments and 10 ms duration) were recorded. Quantification of input resistance of cells was derived via Ohm’s law based on the slope of voltage responses measured at each current step. The inter-pulse interval was 100 ms. Recordings were performed with a Multiclamp 700B amplifier (Molecular Devices). Data were digitized at 10 kHz with a DAQ board (National Instruments, USB-6353) and recorded with a custom software developed in C#.NET and VB.NET in the laboratory. Analysis was done using custom software developed in Delphi and Python environments.

### LFP Recordings

Acute slice preparations were gathered at each recording day (6 slices/animal, 450 µm thick) in a pairwise manner from control and microglia depleted animals while using the same solutions and equipment. The slice preparation sequence was alternated throughout the recording days between the two groups, as well as the chambers that were used for the incubation process, to minimize artefacts that might have been introduced by variance in slice preparation or incubation quality. After at least 1 hour of incubation, slices from both conditions were transferred together in a pairwise manner to a dual perfusion system recording chamber (Hájos and Mody, 2009), and measured simultaneously via performing local field potential (LFP) recordings. In this design, the slices were placed on a metal mesh, and two separate fluid inlets allowed ACSF to flow both above and below the slices at a rate of 3–3.5 ml/min for each flow channel at 33 –34°C (Supertech Instruments). The position of slices from the two conditions were also alternated in the recording chamber between subsequent measurements. Standard patch pipettes filled with ACSF were used for LFP recordings. Recording pipettes were positioned at the same depth (∼80-100 µm below the surface) and in the same region (pyramidal layer of CA3) in both conditions. ACSF containing pipette resistances were 3–6 MΩ. Recordings were performed with a Multiclamp 700B amplifier (Molecular Devices). Data were digitized at 10 kHz with a DAQ board (National Instruments, USB-6353) and recorded with software developed in C#.NET and VB.NET in the laboratory.

### Digital signal processing and analysis

All data were processed and analyzed off-line using self-developed programs written in Delphi 6.0 by A.I.G. and Python 2.7.0 by D.S. Signals were filtered with a two-way RC filter to reserve phase. SWRs were pre-detected on 30 Hz low-pass-filtered field recordings using a threshold value of 2-3 times the SD of the signal. Recordings were considered to not contain SWRs if 2 times the SD of the signal did not result in detectable events. All automatic detection steps were supervised in each recording. The predetected SWRs were then analyzed using a program that measured various SWR features and eliminates recording artefacts similar to SWRs. Namely, on the low-pass-filtered signal, the program measured: peak amplitude of sharp waves (SWR amplitude); inter-sharp wave interval (SWR rate). On a ripple bandpass-filtered trace (200 ±30 Hz), the program also detected the time of negative ripple peaks. Based on this, we identified the ripple cycle closest to the SWR peak and used its negative peak as triggering event for averages to preserve ripple phase. Taking the absolute value of the ripple bandpassed signal and low pass filtering it, we calculated the ripple power peak (Ripple amplitude). After detection, ∼100 consecutive events were selected for quantification, where highest values were measured along the whole recording.

### Quantification and statistical analysis

All quantitative assessments were performed in a blind manner. Sample size was determined based on sample size calculations performed in our previous experiments using similar models. Data were sampled in a systematic random manner. Experiments were replicated by using multiple animals for slice preparations or histology (biological replicates), and pooled results from experiments were presented in the Figs. Exclusion criteria were pre-established for quality of acute slices and immunostainings. No samples had been excluded in the present paper. Normality of the data was tested via Shapiro-Wilk test, to decide whether to use parametric or non-parametric tests during further analysis. In the case of two independent groups, Student’s t-test, or Mann-Whitney U-test, for three or more independent groups Kruskal-Wallis test with Dunn’s multiple comparison test was applied. Statistical tests were conducted in custom or self-developed programs in Python environment 2.7.0 by D.S. In this study, data are presented as mean±SEM or in median-Q1-Q3 format, p<0.05 was considered statistically significant.

## Acknowledgements

We thank László Barna, Pál Vági and the Nikon Imaging Center at the Institute of Experimental Medicine (IEM) for kindly providing microscopy support. We thank Zsolt Kohus for methodological concepts and preliminary electrophysiological data. We are also grateful to Norbert Hájos (IEM), Zoltán Nusser (IEM), and Dániel Schlingloff (IEM) for their support and useful comments.

## Author contributions

Experimental design and overall concept, A.D., C.C., A.G., P.B.; Methodology, C.C., P.B., B.P., Z.K., E.S., H.B., A. A., I.M., X.W., K.H., Y.W., Z.W., M.J., Y.L; Formal Analysis C.C., P.B., B.P., Z.K.; Investigation, P.B., C.C., B.P., E.S., Z.K., A.K., M.N; Resources, A.D., A.G. Writing – Original Draft, P.B., C.C., A.D., Z.K.; Editing, P.B., C.C., B.P., A.D., Z.K., H.B., K.A. I.M., X.W.; Visualization P.B., C.C., B.P., Z.K. Supervision A.D. and A.G.; Project Administration A.D.; Funding Acquisition, A.D.

## Ethics statement

All experiments were approved by the Ethical Committee for Animal Research at the Institute of Experimental Medicine, Hungarian Academy of Sciences, and conformed to Hungarian (1998/XXVIII Law on Animal Welfare) and European Communities Council Directive recommendations for the care and use of laboratory animals (2010/63/EU) (license number PE/EA/2552-6/2016; PE/EA/254-7/2019).

## Supplementary video captions

**Suppl. Video 1.** Time lapse XY (upper) and YZ (lower) views (maximum intensity projections) of a CX3CR1^+/GFP^ microglia moving towards the surface (dotted line) in an acute brain slice. The cell body positions are indicated with coloured dots. Note the extensive process outgrowth and cell body translocation. Scale bar: 10um.

**Suppl. Video 2.** Time lapse volume view (from YZ view) of a CX3CR1^+/GFP^ microglia with almost stationery cell body, showing still strong process redistribution in an acute brain slice.

**Suppl. Video 3.** Morphological changes of individual microglia cells within acute brain slices depicted by maximum intensity projections of XY views. The video starts 1h after slice preparation. Scale bar: 10um.

**Suppl. Video 4.** Morphological differences of individual microglial cells at different timepoints during incubation within acute brain slices depicted by 3D reconstructions via morphological analysis tool (Heindl et al., 2018).

**Suppl. Video 5.** ATP dynamics throughout the 5 hours period after slice preparation recorded from layers of the cortex and the hippocampus

